# Heterotypic Protein Interactions Modulate the Condensate Dynamics and Aggregation of α-Synuclein

**DOI:** 10.1101/2025.07.10.664163

**Authors:** Sumangal Roychowdhury, Souradip Paul, Kandarp A. Sojitra, Sounak Bhattacharya, Abhishek Mazumder, Jeetain Mittal, Krishnananda Chattopadhyay

**Affiliations:** Structural Biology & Bio-Informatics Division, CSIR-Indian Institute of Chemical Biology, 4, Raja S.C. Mallick Road, Kolkata 700032, India; Academy of Scientific and Innovative Research (AcSIR), CSIR-Human Resource Development Centre, Ghaziabad 201002, India; Artie McFerrin Department of Chemical Engineering, Texas A&M University, College Station, TX 77843, USA; Central Instrumentation Facility, CSIR-Indian Institute of Chemical Biology, 4, Raja S.C. Mallick Road, Kolkata 700032, India; Department of Chemistry, Texas A&M University, College Station, TX 77843, USA; Interdisciplinary Graduate Program in Genetics and Genomics, Texas A&M University, College Station, TX 77843, USA

## Abstract

Multicomponent biomolecular condensates formed by diverse multivalent proteins underlie numerous cellular processes, from ribosome biogenesis to stress response regulation, and have recently been implicated in disease-related protein aggregation. Understanding how heterotypic protein interactions modulate condensate dynamics and transitions to amyloid states remains a major challenge. Here, we combine fluorescence -based ensemble and single-molecule approaches, molecular simulations, and systematic domain deletions to investigate how charge patterning and domain structure influence co-condensation between α-synuclein, a disordered neuronal protein linked to Parkinson’s disease, and the SARS-CoV-2 nucleocapsid protein (NP), a structured viral RNA-binding protein. We find that co-condensation is driven by multivalent electrostatic interactions and occurs when the heterotypic affinity exceeds a threshold, resulting in restricted protein dynamics and altered condensate material properties. These changes promote the formation of dense amyloid fibrils, as confirmed by atomic force microscopy and fluorescence assays. Our results elucidate how heterotypic interactions within multiphasic condensates can modulate phase behavior and aggregation, providing insight into broader mechanisms linking condensate dysregulation with neurodegenerative diseases.

## Introduction

In recent years, the formation of biomolecular condensates or membraneless organelles (MLOs) via liquid-liquid phase separation (LLPS) has emerged as the process underlying many biological functions^7,8^. These condensates arise from demixing of proteins and nucleic acids into two phases: a condensed phase with high concentration and a dilute phase with low concentration of biomolecules^9^. Previous studies have shown that proteins with intrinsically disordered regions (IDRs) can undergo phase separation by promoting multivalent interactions between polypeptide chains^10,11,12,13^. In addition, several globular or well-folded proteins can also phase separate^14^, due to either molecular crowding (BSA, lysozyme)^15^ or local unfolding of the protein chain (SOD1)^16^. These condensed phases can selectively recruit other biomolecules such as proteins and nucleic acids to form co-condensates. Recent developments suggest that a balance between homotypic and heterotypic interactions in scaffold and client molecules can guide the co-partitioning of IDRs^17,18^. Factors such as domain architecture, amino acid composition, oligomerization, and the influence of binding partners can modulate these interactions, forming co-mixed or multiphasic condensates^19^. However, the mechanisms governing co-condensation in multidomain proteins remain unclear, limiting our understanding of their role in disease.

Here, we studied co-condensation between a-synuclein (α-syn) and SARS-CoV-2 nucleocapsid protein (NP) under *in vitro* condition. Numerous studies have shown that α-syn forms biomolecular condensates in the presence of macromolecular crowders or inside neuroblastoma cells under stress^20,21^. NP can also form liquid droplets in the presence of nucleic acids (genomic RNA) or form co-condensates with hnRNPA2, TDP-43, and FUS ^22,23,24,25^. A recent study has proposed that NP may accelerate neurodegeneration by promoting the formation of amyloid fibrils composed of the PD-related protein α-syn^5,26^. It is now believed that nucleation occurs through condensate formation, which provides a suitable reaction crucible facilitating interactions between transiently formed higher-order species or oligomers^12,27,28,29^. In this scenario, the involvement of other client proteins or small molecules can accelerate or induce phase separation and aggregation by acting as binding partners or by altering the conformation of scaffold proteins. Given that α-syn aggregation is nucleation-driven and often initiated within condensates, understanding how NP influences this process is critical.

In this study, we show that full-length NP can enhance the condensation of α-syn. Using Fluorescence Correlation Spectroscopy (FCS), Fluorescence Recovery After Photobleaching (FRAP), and Time-domain Fluorescence Lifetime Imaging (TD-FLIM), we examined the dynamics of α-syn within the co-condensates (composed of α-Syn and NP) and inside α-syn only condensates. Coarse-grained (CG) simulations revealed that interactions are mediated by oppositely charged blocks in the C-terminal domain (CTD) of α-syn and different domains of NP. To further investigate the role of individual domains of the two proteins in their co-condensation, we purified domain deleted variants of α-Syn and NP and used confocal microscopy to examine their phase behaviors, which ranged from co-mixed to core-shell architectures. Binding affinity measurements using fluorescence anisotropy enabled us to establish a correlative relationship between the partition coefficient of NP inside condensates and the heterotypic binding affinity (K_d_), suggesting that heterotypic interactions drive co-condensation when this threshold is surpassed. The temporal maturation of the co-condensates showed increasing partitioning of an amyloid reporter, Thioflavin-T (ThT), indicating the formation of cross β-sheet structures inside the co-condensates. Atomic Force Microscopy (AFM) combined with ThT fluorescence confirmed a higher aggregation tendency and dense amyloid fibrils of α-syn in the presence of NP. Overall, our findings uncover how SARS-CoV-2 NP drives co-condensation with α-syn and accelerates amyloid formation, shedding new light on the interplay between viral infection and neurodegenerative disorder and offering potential avenues for therapeutic intervention.

## Results

### NP enhances the phase separation of α-syn by forming co-condensates

The protein α-syn is intrinsically disordered, featuring an amphipathic N-terminal domain (NTD), a hydrophobic non-amyloid component (NAC) domain in the middle, and a predominantly disordered acidic CTD **(Figure 1A, S1A)**^30^. In contrast, NP is a multi-domain protein composed of three predicted intrinsically disordered regions (the NTD known as N_IDR_, the Serine/Arginine-rich linker region, and the CTD known as C_IDR_), an RNA-binding NTD, and a dimerizing CTD **(Figure 1A, S1B)**^22^. At pH 7.4, α-syn’s primary sequence carries a net negative charge of -9, whereas NP corresponds to a net positive charge of +24 **(Figure 1A)**. Prior work has shown that oppositely charged polyelectrolytes can lead to complex coacervation through heterotypic charge attractions, a process observed in other biological systems such as ProTα/H1, PrP/α-syn, etc^31,32^.

**Figure 1:**
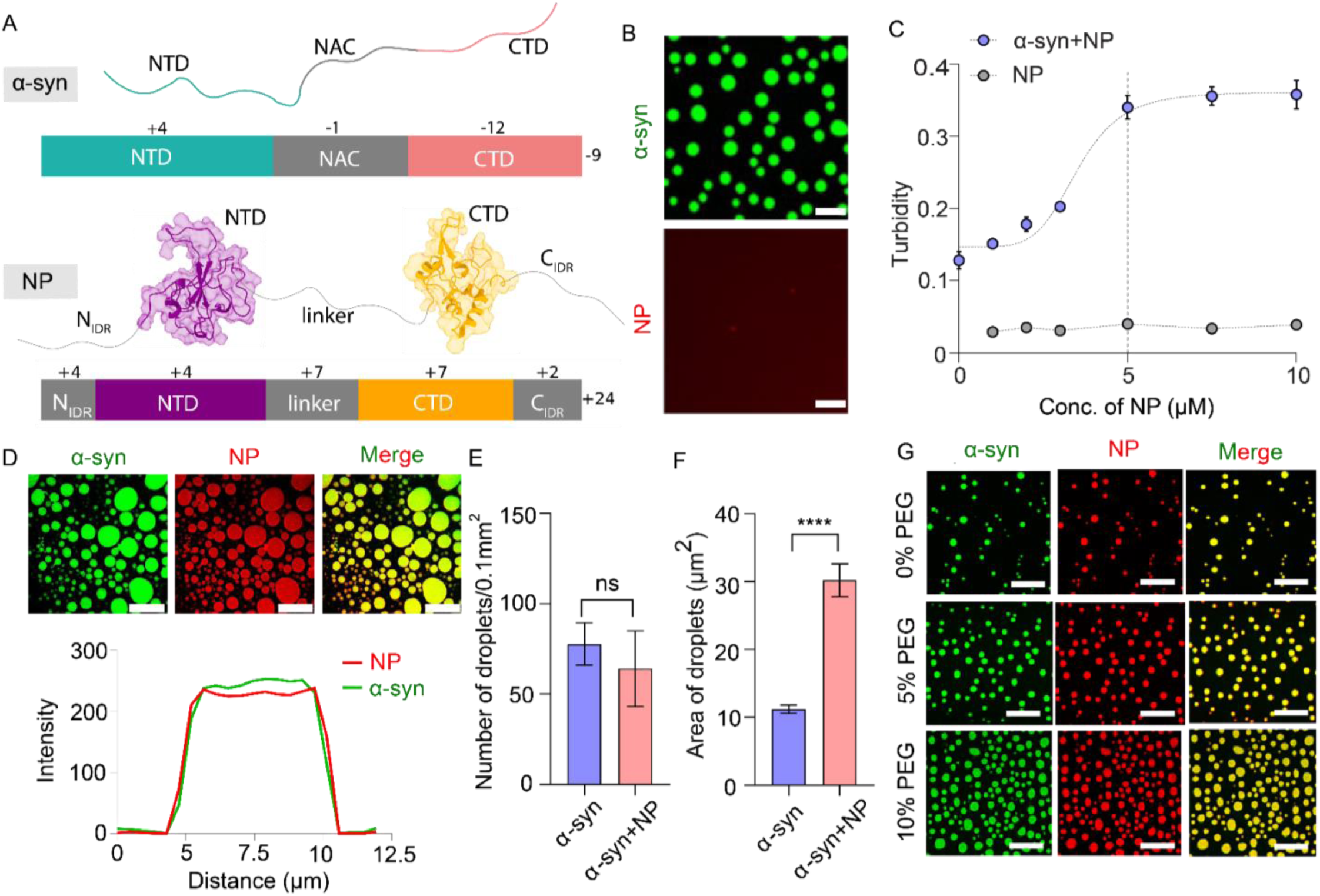
Co-condensation of α-syn and NP. (A) (Upper) Cartoon representation of disordered α-syn and its domain structure displayed below it. (Lower) Cartoon representation of NP (PDB ID: NTD- 7ACT; CTD- 6ZCO) and its domain structure displayed below it. Different colors indicate different domains of the proteins. The net charge of each domain and entire protein is displayed above and to the right of the domain architecture. (B) (Upper) Condensates of α-syn after ∼ 4 hours of incubation at 37τC and 180 rpm (100µM unlabeled protein was doped with 1% Alexa-488 maleimide-labeled protein, 10% PEG-8k, 400mM MgSO_4_). (Lower) No droplets were observed for NP (5µM unlabeled protein was doped with 1% Alexa-647 NHS ester labeled protein, 10% PEG-8k, 400mM MgSO_4_). Three independent experiments were performed (n=3). Scale bar: 10µm (C) Turbidity (light scattering measured at 350 nm) plot of α-syn (100µM unlabeled α-syn incubated with 10% PEG-8k, 400mM MgSO_4_ at 37τC, 180 rpm for 4 hours) and α-syn in the presence of NP (100µM unlabeled α-syn and 5µM unlabeled NP were incubated with 10% PEG-8k, 400mM MgSO_4_ at 37τC, 180 rpm for 4 hours). Data are shown as mean± SD (n=5). (D) (Upper) Confocal microscopic images of co-condensates of α-syn doped with 1% Alexa-488-labeled α-syn and NP doped with 1% Alexa-647-labeled NP (Molar ratio of α-syn and NP is 20:1) in the presence of 10% PEG-8k, 400mM MgSO_4_ after ∼ 4 hours of incubation at 37τC and 180 rpm. Scale bar: 10µm. Three independent experiments were performed (n=3). (Lower) Line intensity vs distance plot of ɑ-syn and NP co-condensate showing complete co-localization of ɑ-syn and NP droplet. (E) Plot depicting the number of droplets observed in confocal microscopic images for homotypic α-syn droplets and heterotypic co-condensates of α-syn and NP. Data are shown as mean± SEM (n=5). “ns” indicates non-significant. P values were determined by non-parametric t-test. (F) Plot depicting the area of droplets observed in confocal microscopic images for homotypic α-syn droplets and heterotypic co-condensates of α-syn and NP. Data are shown as mean± SEM (n=5). **** indicates P value < 0.0001. P values were determined by non-parametric t-test. (G) Confocal microscopic images showing the effect of crowding agent (PEG-8k) on the co-condensation of Alexa-488 labeled α-syn and Alexa-647 labeled NP. The percentage of PEG-8k added is indicated on the left of each figure. Scale bar: 20µm. Three independent experiments were performed (n=3).

Before probing binary heterotypic phase separation, we first experimentally investigated the homotypic phase separation of both proteins individually in vitro by incubating α-syn and NP separately at pH 7.4 and 37°C under constant stirring (180 rpm). We also used 10% polyethylene glycol (PEG) as a crowding agent to mimic the cellular environment. We observed that α-syn formed spherical droplets (∼1-3 µm in diameter) at a concentration of 100 μM, whereas NP did not form any homotypic droplets at concentrations up to 15 μM **(Figure 1B, S1C)**. To further examine the effect of NP on α-syn condensation, we varied NP concentrations while maintaining α- syn at 100 μM (the concentration used for homotypic condensation) and assessed the optical turbidity via light scattering measurements at 350 nm. The NP concentration dependence shows a significant increase in light scattering, saturating at ∼5 μM NP **(Figure 1C)**. Overall, these results demonstrate that mixing NP with α-syn significantly enhances phase separation compared to the homotypic phase separation of α-syn alone. For all subsequent experiments, we used a stoichiometric mixture of 100 μM α-Syn and 5 μM NP to induce heterotypic phase separation.

Next, we performed confocal microscopy to directly visualize the mesoscopic droplets. To visualize liquid droplets under confocal microscopy, we labeled α-Syn using Alexa488 c5 Maleimide at a cysteine residue, which we inserted at the 132^nd^ position in the protein. In contrast, NP was labeled at the N-terminal using Alexa 647 NHS ester. For all microscopy experiments, ∼1% fluorescently labeled proteins were used to dope higher concentrations (100 μM α-Syn and 5 μM NP) of unlabeled proteins **(Figure 1D)**. It is known that salt concentration modulates complex coacervation by screening electrostatic interactions and through ion-specific effects, as described by the Hofmeister series. Certain salts can promote or inhibit phase separation depending on their identity and concentration.^33,34,35,36^ Keeping these effects in mind, we next examined how ionic strength influences the heterotypic phase separation of α-syn and NP using varying concentrations of NaCl, MgSO_4_, Na_2_SO_4_, and MgCl_2_ **(Figure S1D-G).** By monitoring turbidity over time **(Figure S1H)**, we optimized an incubation period of 240 mins for these measurements. Using microscopy, we observed that at relatively lower NaCl concentrations (200 mM and 400 mM), the droplets appeared sticky and clustered **(Figure S1D, S1I)**. However, in the presence of 800 mM NaCl, these two proteins were found co-localized within well-defined condensates. In contrast, in the presence of MgSO₄, Na₂SO₄, and MgCl₂, high concentrations led to sticky and clustered droplets, whereas lower concentrations significantly improved droplet morphology **(Figure S1E–G, S1I)**. **Figure S1J** additionally shows that a solution condition with ionic strength between 0.6 and 1.6 may be optimal for the heterotypic phase separation of these proteins. Close inspection of the microscopy data **(Figure S1E, middle panel and S1K-L)** reveals that 400mM MgSO_4_ resulted in the highest number of heterotypic droplets and the largest total droplet area. This condition was therefore used for all subsequent experiments.

When we compared homotypic (α-syn-only) and heterotypic (α-syn/NP) droplets, we noted that the number of the droplets were similar, but the droplet areas were significantly larger in the heterotypic droplets **(Figure 1E-F).** Finally, because the presence of a crowding agent (like PEG) can facilitate both heterotypic and homotypic phase separation^42,43^, we tested different concentrations of PEG-8000 (0%, 5%, and 10% as shown in **Figure S1M, 1G**). Based on the data, we used 10% PEG-8000 in all subsequent experiments. Under these optimized conditions, the droplets showed rapid fusion within few seconds, indicating liquid-like behavior (Supplementary video 1). Also, the data outlined in this section clearly show that NP promotes α-syn phase separation, with condensate formation influenced by ionic strength and molecular crowding, thereby providing a basis for further investigation.

### α-syn shows slower dynamics in heterotypic co-condensates

Recently, it has been shown that in α-syn, the hydrophobic NAC region plays a key role in regulating phase separation through intermolecular NAC-NAC interactions^44^. In the presence of salt, long-range interactions between the NTD and the CTD are screened^45^, exposing the hydrophobic NAC domains, which enhances phase separation and aggregation. To probe the overall hydrophobic environment of both homotypic and heterotypic droplets formed by α-syn and NP, we used an environment-sensitive solvatochromic dye Merocyanine 540 (MC-540). It has been reported that the fluorescence intensity of MC-540 is enhanced when it enters a hydrophobic environment^46^ **(Figure S2A).** This dye has been used to detect hydrophobic microenvironments and spatial inhomogeneities within A1-LCD condensates^47^. In our study, we prepared homotypic (α-syn only) and heterotypic (α-syn/NP) droplets doped with 50 µM MC-540. Confocal microscopy and dye partitioning analyses revealed that the partition coefficient of MC-540 was significantly higher in the co-condensates of α-syn and NP compared to those of α-syn droplets **(Figure S2B-C)**, indicating an increased hydrophobic environment inside the co-condensates.

Intrinsically disordered proteins (IDPs) exhibit high flexibility and frequent local rearrangements even in densely packed environments^48,49,50,51^. This means that while the overall structure may appear stable, individual protein molecules are in constant motion, interacting within a dynamic network and undergoing conformational changes on short timescales^11,52,53^. Previous literature has shown that the presence of a client molecule (which can be a protein or a nucleic acid) can affect the dynamic behavior of the scaffold proteins inside the condensates^31^.

Here, we studied the diffusional dynamics of α-syn in the presence of NP using fluorescence correlation spectroscopy (FCS). FCS monitors fluorescence intensity fluctuations of a probe within a small confocal volume (∼femtoliter scale) **(Figure 2A)**. For these studies, we doped the unlabeled solution of both α-syn alone and a mixed solution of α-syn and NP with 15nM Alexa-488-labeled α-syn. We used point-FCS technique^16^ to measure α-syn diffusion in both dilute and dense phases (inside the droplets). The normalized single-color autocorrelation functions obtained from FCS were fit to a suitable model, and diffusion coefficient values were determined using previously described methods^16,54^. When we imaged the droplets, we found that heterotypic droplets were brighter than homotypic droplets **(Figure S3A)**. While this suggests a relatively higher number of α-syn molecules inside heterotypic droplets, some caution is warranted, as (1) only a small percentage of α-syn molecules are labeled and hence probed, which may not necessarily represent the overall population; and (2) there could be a change in the fluorescence lifetime of the probes affecting intensity values.

**Figure 2:**
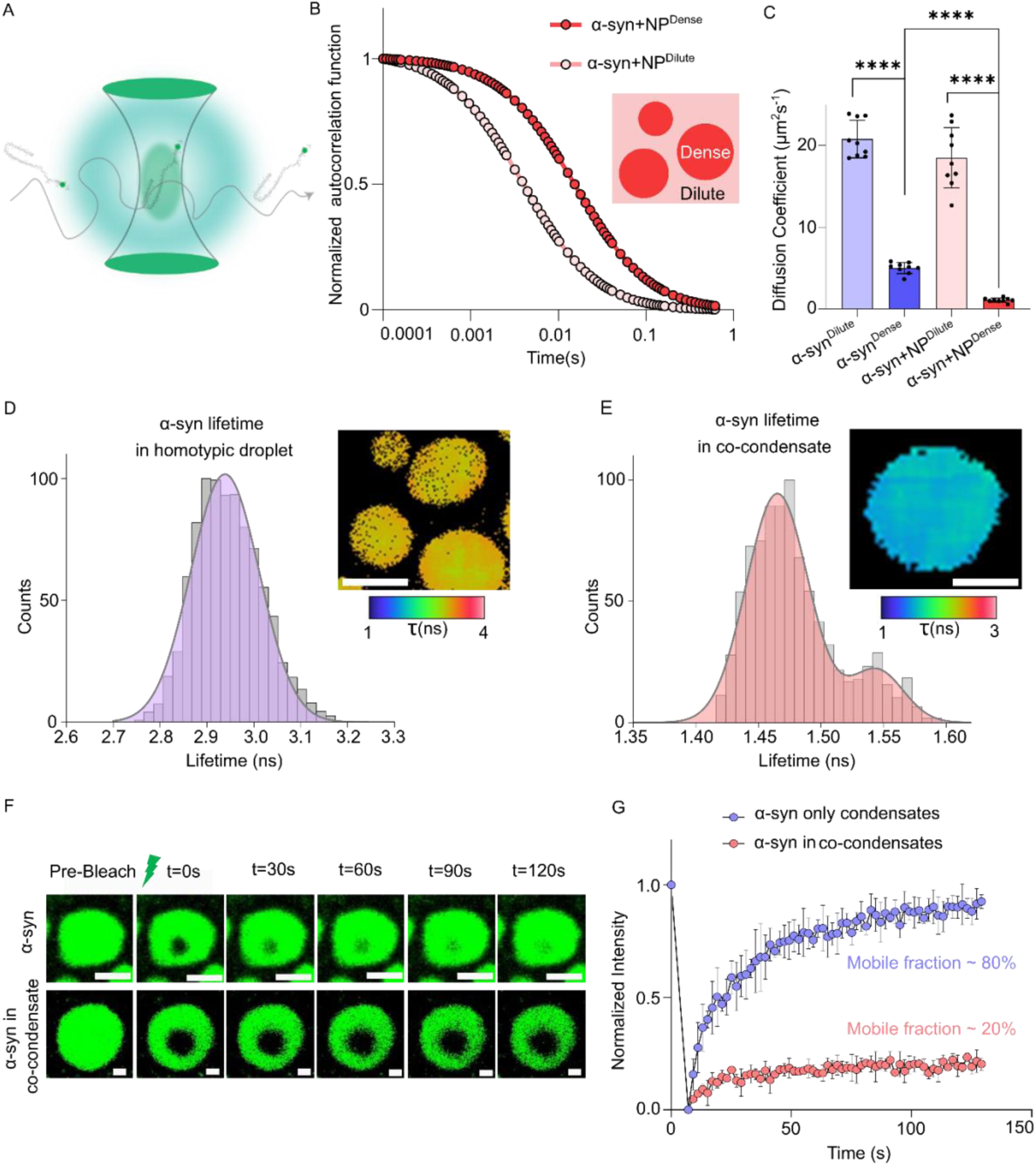
Reduced dynamics of α-syn in heterotypic condensates. (A) Schematic representation of single molecule FCS measurement, showing a labeled α-syn molecule diffusing through the confocal volume. (B) Amplitude normalized autocorrelation function obtained from FCS of α-syn in the co-condensates. Intense colors indicate the dense phase of the co-condensates, and lighter variants indicate the dilute phase of the co-condensates, as indicated in the schematic shown in the inset. The shift in the autocorrelation function indicates a longer diffusion time in the dense phase compared to the dilute phase. (C) Plot depicting a comparison of diffusion coefficients obtained from FCS measurements of dense and dilute phases of homotypic and heterotypic droplets. Data are shown as mean± SD (n=9). **** indicates P value < 0.0001. Statistical significance was established by a non-parametric t-test. (D) Histogram plot of lifetimes obtained from TD-FLIM measurements of homotypic droplets of α-syn. The inset shows the representative lifetime images of homotypic α-syn condensates. Scale bar: 10µm. The color bar under the image indicates the range of lifetime values in the legend. (E) Histogram plot of lifetimes obtained from TD-FLIM measurements of heterotypic droplets of α-syn and NP. The inset shows the representative lifetime images of α-syn-NP co-condensates. Scale bar: 10µm. The color bar under the image indicates the range of lifetime values in the legend. (F) Representative droplet image at indicated time points of FRAP measurements for homotypic α-syn condensates and heterotypic droplets of α-syn and NP. Scale bar: 1.5µm (G) FRAP kinetics curve for α-syn in homotypic α-syn condensates (violet) and α-syn in heterotypic droplets of α-syn and NP (pink). The mobile fractions for α-syn in homotypic α-syn condensates were ∼ 80% and α-syn in heterotypic droplets of α-syn and NP were ∼ 20%, indicating arrested dynamics of α-syn in the co-condensates. Data is shown as mean± SD (n=5) for five independent experiments.

We also observed a large shift of FCS autocorrelation traces in the dense phase compared with the dilute phase for both homotypic (**Figure S3B**) and heterotypic droplets (**Figure 2B**). In homotypic condensates of α-syn, the diffusion coefficient in the dense phase was ∼4 times lower than that in the dilute phase. In contrast, within heterotypic co-condensates, we detected a much larger decrease in the diffusion coefficient of α-syn in the dense phase (∼17 times) compared to homotypic condensates **(Figure 2C)**. This observation suggests that heterotypic droplets are more densely packed than homotypic droplets. We note that the diffusion coefficient values in the dilute phase were nearly the same for both homotypic and heterotypic condensates, and the ∼10% decrease was expected, as the molecular mass of the α-syn-NP complex is larger than α-syn alone. Notably, this decrease in the diffusion coefficient in the dilute phase is similar to what we measured in the solution phase (without incubation for droplet formation) for α-syn in the presence and absence of NP **(Figure S3C)**. The corresponding FCS autocorrelation traces are shown in **(Figure S3D-E).**

We then used time-domain fluorescence lifetime imaging (TD-FLIM) to investigate the spatial changes in the protein’s microenvironment inside the droplets. Notably, the lifetime of the fluorophore is independent of protein concentration. The analysis of TD-FLIM revealed that the lifetime distribution of Alexa-488-maleimide-tagged α- syn exhibited a broad peak with a mean lifetime of 2.93 ns **(Figure 2D)**, which is slightly lower than that of monomeric α-syn measured in dilute solution (3.44 ns). This type of broad lifetime distribution is typical for other disordered proteins like ProTα, FUS, and unfolded ubiquitin, as shown earlier^31,55–57^. However, inside the co-condensates, we observed a bimodal distribution with significantly lower α-syn lifetime values (1.47 ns and 1.55 ns) **(Figure 2E)**. The representative time-correlated single-photon counting (TCSPC) decay curves for homotypic and heterotypic droplets are shown in **Figure S3F**. The presence of a bimodal distribution presumably suggests a higher degree of heterogeneity inside the co-condensate compared to α-syn-only condensates. In addition, significantly lower fluorescence lifetime inside the co-condensates indicates a more rigid environment in the presence of NP. However, the lifetime inside the co-condensates is lower, it remains higher than the lifetime value observed for α-syn aggregates (∼1 ns) reported previously in vitro and in model organisms^58^.

We further characterized the diffusional dynamics inside the condensates using Fluorescence Recovery After Photobleaching (FRAP). Homotypic α-syn droplets showed ∼80% recovery, indicating fast translational diffusion of the protein inside the droplets **(Figure 2F-G)**. α-syn exhibited much slower recovery (∼20%) in the co-condensates, indicating arrested protein dynamics **(Figure 2F-G)**. In contrast, when we measured FRAP of NP inside the co-condensates, it showed an even slower recovery of ∼5%, suggesting hindered motion relative to α-syn **(Figure S3G)**. This type of altered dynamics was also observed earlier in the heterotypic co-condensation of nucleolar protein Nucleophosmin1 (NPM1) with non-ribosomal nucleolar protein Surfeit locus protein 6 (SURF6)^59^. In summary, all three techniques - FCS, FRAP, and FLIM revealed a more conformationally constrained environment for α-syn in the dense phase of heterotypic condensates compared to homotypic droplets.

### Molecular simulations & domain deletion experiments provide insights into the formation of co-condensates

Our experimental results clearly show that NP enhances α-syn condensation. To understand the interactions stabilizing their co-condensation, we conducted coexistence CG simulations **(Figure S4A)**. Our analysis revealed that α-syn CTD forms substantial contacts with both the folded and disordered domains of NP **(Figure 3A-B)**. Given their opposite net charges **(Figure 1A)**, charge complementarity could be a factor driving the co-condensation. Previous studies have shown that segregated charges in IDRs influence compaction and promote homotypic condensation, while local charged blocks can also regulate client partitioning within scaffold IDRs^60,61,62,63,64^. A closer look at the charge distribution showed that α-syn has more negative charges at the C-terminus, whereas NP exhibits local charge clustering **(Figure 3B, S4B)**. To investigate this further, we identified the positively and negatively charged blocks, as described by Lyons et al.^60^. We observed two negatively charged blocks at the CTD of α-syn and three positively charged blocks in NP, one each in the N_IDR_, NTD, and CTD **(Figure 3B)**. Based on these observations, we hypothesized that the oppositely charged blocks could drive co-condensation. Specifically, we posited that: (a) deleting the NTD or CTD in NP would decrease co-partitioning, by removing positive blocks that interact with the negative block in α-syn; (b) similarly, deleting the CTD in α-syn would remove the negative block, thereby decreasing the partition coefficient; (c) however, deleting the NTD in α-syn would likely not decrease the partition coefficient because the negative block remains, and thus maintain strong interaction with NP.

**Figure 3:**
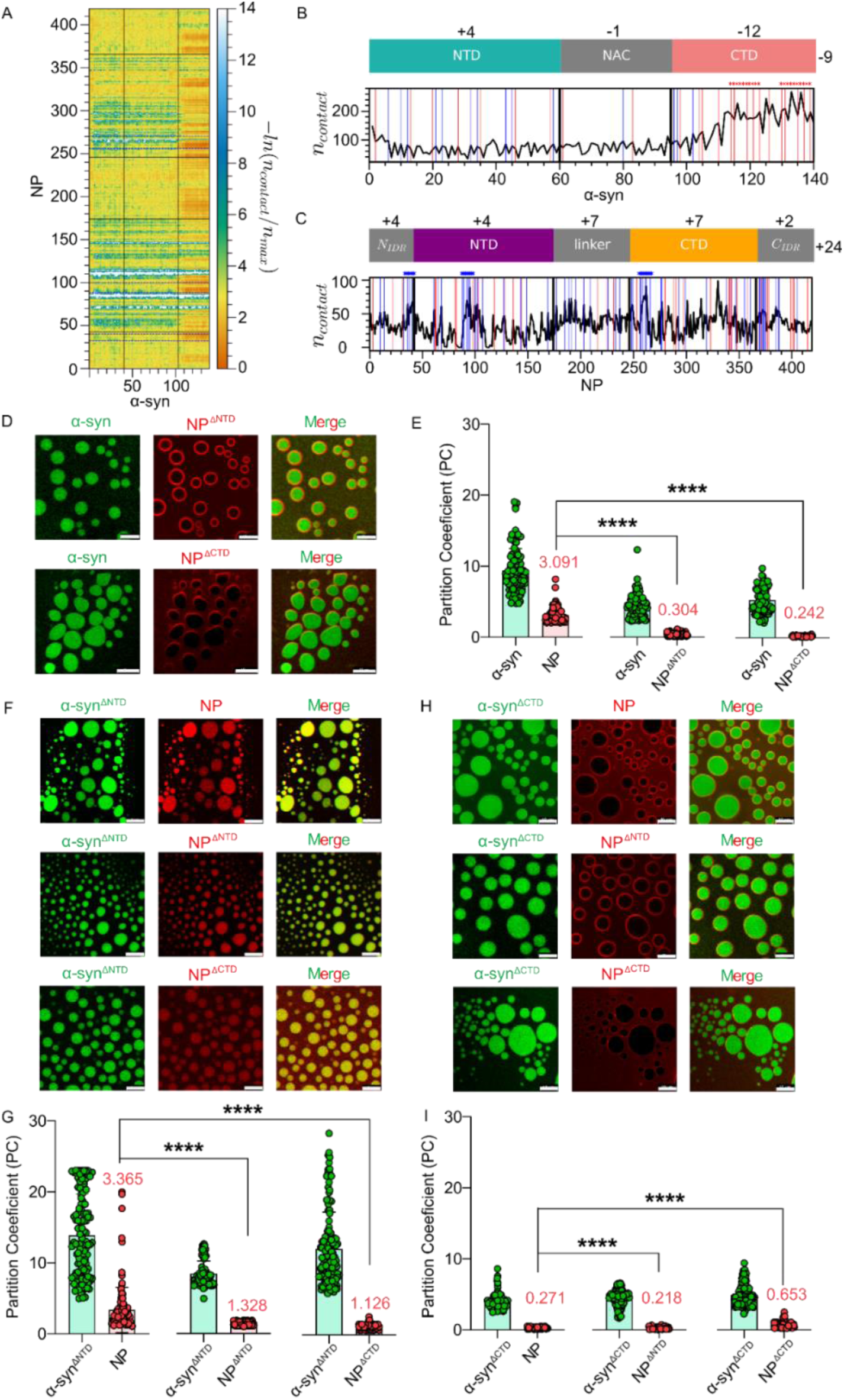
Domain-deleted mutants of α-syn and NP showed different spatial arrangements. (A) Pairwise residue index contact map between α-syn and NP obtained from CG co-existence simulations. (B) (Upper) Domain architecture of α-syn with net charge distribution. (Lower) Per residue contacts formed by α-syn with NP in CG co-existence simulations, with negatively charged blocks identified using a 10-residue sliding window (as described by Lyons et al.)^58^ (C) (Upper) Domain architecture of NP with net charge distribution. (Lower) Per residue contacts formed by NP with α-syn in CG co-existence simulations, with positively charged blocks computed as in (B). (D) Confocal images of co-condensates formed by Alexa-488-labeled α-syn and Alexa-647-labeled NP with N-terminal (Upper) or C-terminal (Lower) deletion. Core-shell droplets were observed, with NP primarily in the shell and α-syn in the core. Scale bar: 10 µm. (E) Plot of the partition coefficient of individual proteins in the co-condensates formed involving α-syn and NP, or their domain-deleted mutants, as indicated in the x-axis. Data are shown as mean± SEM (n=5) for five independent experiments. **** indicates P value < 0.0001. Statistical significance was established by a non-parametric t-test. (F) Confocal images of co-condensates involving N-terminal deleted α-syn and NP (Upper), N-terminal deleted α-syn with N-terminal deleted NP (middle), and N-terminal deleted α-syn with C-terminal deleted NP (Lower). Complete co-localized droplets were observed in all cases. Scale bar: 10 µm. (G) Plot of the partition coefficient of individual proteins in the co-condensates formed involving N-terminal deleted α-syn and NP or their domain-deleted mutants, as indicated in the x-axis. Data are shown as mean± SEM (n=5) for five independent experiments. **** indicates P value < 0.0001.(H) Confocal images of co-condensates involving C-terminal deleted α-syn and NP (Upper), C-terminal deleted α-syn with N-terminal deleted NP (Middle), and C-terminal deleted α-syn with C-terminal deleted NP (Lower). Core-shell droplets were observed, with NP primarily in the shell and α-syn in the core. Scale bar: 10 µm (I) Plot of the partition coefficient of individual proteins in the co-condensates formed involving C-terminal deleted α-syn and NP or their domain-deleted mutants as indicated in the x-axis. Data are shown as mean± SEM (n=5) for five independent experiments. **** indicates P value < 0.0001. All confocal microscopy experiments were performed five times (n=5) with similar results.

To experimentally validate these predictions, we systematically deleted different domains of both α-syn and NP and performed microscopy experiments under the previously specified conditions. In the first set of experiments, we used the full-length scaffold protein, α-syn, and deleted the NTD and CTD domains of NP. We co-incubated α-syn and NP^ΔNTD^ and observed that the NP was excluded from the α-syn droplets and primarily localized at the surface of α-syn in a core-shell-like architecture **(Figure 3D, S4C)**. A similar spatial organization was observed upon deletion of the CTD from NP (NP^ΔCTD^) **(Figure 3D, S4C)**. Quantification of partitioning further suggests that co-localization decreased upon either NTD or CTD deletion in NP **(Figure 3E)**.

In the next set of experiments, we deleted amphipathic NTD of α-syn (α-syn^ΔNTD^) and co-incubated it separately with different NP variants (NP, NP^ΔCTD^, and NP^ΔNTD^). We noted that in all three cases, the proteins were co-localized, similar to the case when both WT variants were used **(Figure 3F, S4D)**. Thus, deletion of the NTD in α-syn did not drastically affect the architecture of the co-condensates. However, the extent of partitioning of domain-deleted NP decreased upon NTD deletion in α-syn compared to the WT **(Figure 3G)**. Finally, we turned to our next variant, i.e., CTD-deleted α-syn (α-syn^ΔCTD^). When we incubated α-syn^ΔCTD^ with NP variants (NP, NP^ΔCTD^, and NP^ΔNTD^), we again observed that NP was excluded from the α-syn droplets, forming a core-shell structure **(Figure 3H, S4E)**. The partition coefficient further supported the co-phase behavior observed upon deleting the NTD and CTD of α-syn **(Figure 3G, 3I)**.

In summary, our domain-specific microscopic observations aligned with the predictions from molecular simulations and charged block analysis, revealing that co-phase behavior can be modulated by deleting different domains containing charged blocks of the scaffold and client proteins^60^.

### The binding between α-syn and NP dictates the spatial arrangement of the co-condensates

To quantify the driving force underlying our domain-specific microscopic observations, we determined the binding affinity through a fluorescence-based binding readout, namely fluorescence anisotropy. It is a powerful technique, particularly effective when the fluorescence intensity of a protein labeled with a fluorophore remains unchanged during protein-protein or protein-ligand interactions. We strategically segmented the heterotypic binding assays, similar to our domain-specific confocal experiments. The change in the anisotropy of 100nM labeled α-syn was measured by incrementally increasing the concentration of NP. To determine the heterotypic binding affinity, we fit the data with a one-site binding isotherm **(Figure 4A-C)**.

**Figure 4:**
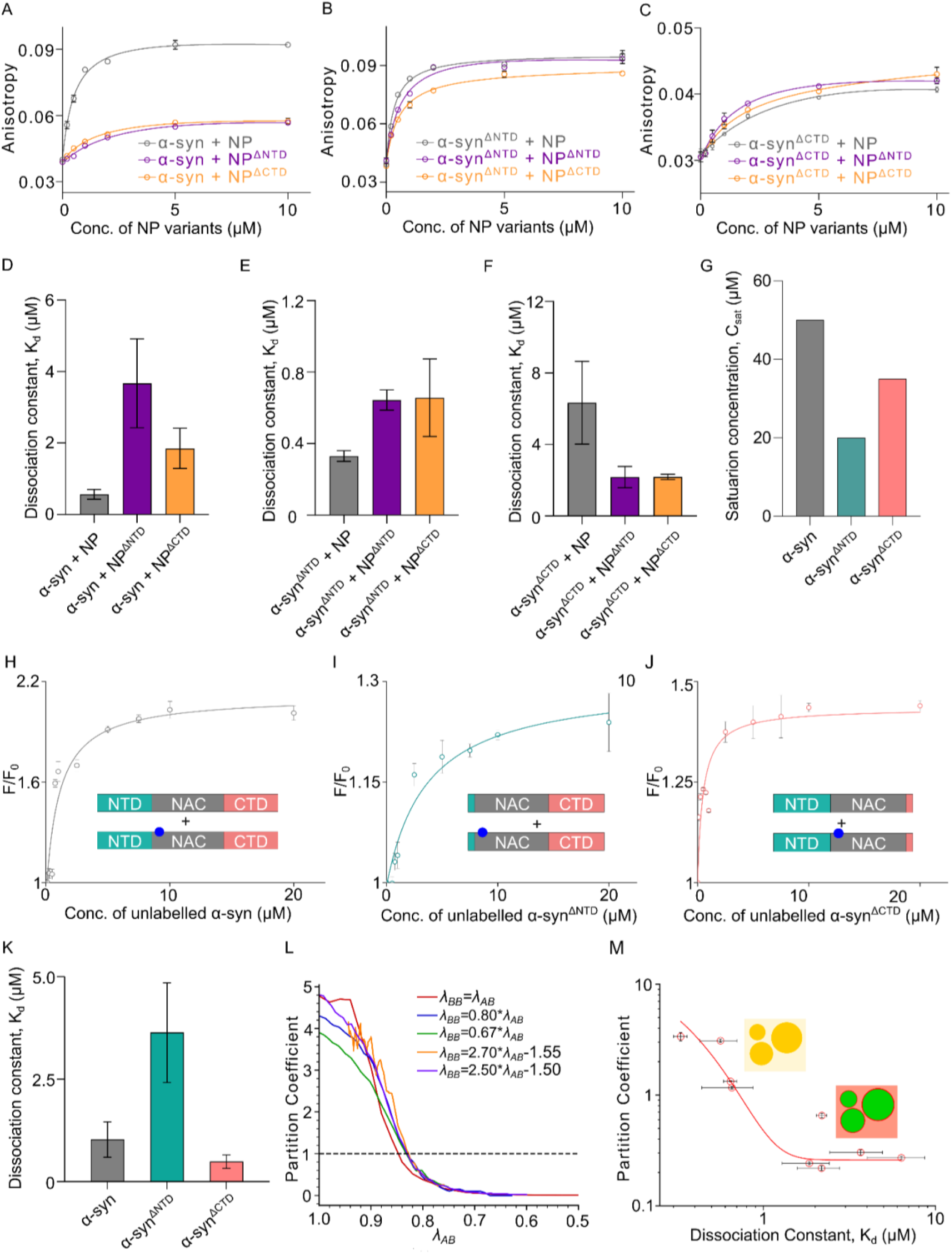
Domain-specific binding interaction dictates the spatial arrangement of co-condensates. (A) Plot depicting anisotropy vs concentration of different variants of NP added to determine binding constants between α-syn and indicated variants of NP using fluorescence anisotropy experiment. Data are shown as mean± SEM for three independent experiments (n=3). (B) Plot depicting the anisotropy vs concentration of different variants of NP added to determine binding constants between N-terminal deleted α-syn and indicated variants of NP using fluorescence anisotropy experiment. Data is shown as mean± SEM for three independent experiments (n=3). (C) Plot depicting the anisotropy vs concentration of different variants of NP added to determine binding constants between C-terminal deleted α-syn and indicated variants of NP using fluorescence anisotropy experiment. Data is shown as mean± SEM for three independent experiments (n=3). (D)-(F) Plots of respective binding constants determined from the fluorescence anisotropy experiments. Data are shown as mean±SEM for three independent experiments (n=3). (G) A plot of respective saturation concentration of different constructs of α-syn as determined from confocal microscopy experiments. (H) Plot showing the normalized fluorescence intensity of Acd-α-syn as a function of the concentration of its unlabeled form to derive a binding isotherm, with the constructs used depicted in the inset. Data are shown as mean± SEM for three independent experiments (n=3). (I) Plot showing the normalized fluorescence intensity of N-terminal deleted Acd-α-syn as a function of the concentration of its unlabeled form to derive a binding isotherm, with the constructs used depicted in the inset. Data is shown as mean± SEM for three independent experiments (n=3). (J) Plot showing the normalized fluorescence intensity of C-terminal deleted Acd-α-syn as a function of the concentration of its unlabeled form to derive a binding isotherm, with the constructs used depicted in the inset. Data is shown as mean± SEM for three independent experiments (n=3). (K) Plot of respective binding constants of homotypic interactions determined from the fluorescence equilibrium stoichiometry titration. Data are shown as mean±SEM for three independent experiments (n=3). (L) Plot depicting variation in partition coefficient as a function of heterotypic interaction strength (λ_AB_) from a minimal polymer model of two polymer chains. The homotypic strength of scaffold polymer is fixed (λ_AA_ = 1) while the variation in homotypic strength of client polymer (λ_BB_) is shown in legend. (M) Plot depicting the partition coefficient of NP constructs in all combinations of heterotypic droplets involving different α-syn and NP variants against their respective dissociation constant as a measure of heterotypic binding strength in log-log scale. The inset scheme illustrates co-localized droplets (partition coefficient above 1) having low heterotypic dissociation constant and multiphasic droplets (partition coefficient below 1) having high heterotypic dissociation constant.

The experiment performed with α-syn and different NP variants (NP, NP^ΔCTD^, and NP^ΔNTD^) revealed that the K_d_ for the WT variant was significantly lower (0.564 ± 0.137 µM) than that of NP^ΔCTD^ (1.848 ± 0.563µM) and NP^ΔNTD^ (3.672 ± 1.246µM) **(Figure 4D)**. Therefore, reduced binding affinity may underlie the expulsion of NP from the co-condensates. When we deleted the NTD of α-syn, the K_d_ remained comparable to that between both WT variants **(Figure 4E)**. These binding affinities also support our confocal data, where we found complete co-localization of these two proteins inside the condensates. Surprisingly, removing the CTD of α-syn (α-syn^ΔCTD^) increased the dissociation constant by ∼12-fold compared to the WT variants. The K_d_ also rose by ∼4- and ∼3.5-fold for NP^ΔNTD^ and NP^ΔCTD^, respectively **(Figure 4F)**. This impaired heterotypic interaction was also reflected in the localization of proteins within co-condensates.

Interestingly, NP^ΔNTD^ and NP^ΔCTD^ variants exhibited co-mixing with α-syn^ΔNTD^, albeit with a lower partition coefficient, even though the positively charged blocks were removed. This suggests that α-syn’s homotypic interactions may still play a role. Previous studies have shown that the homotypic strength of a scaffold protein can influence co-mixing with client proteins, with overall co-mixing governed by an interplay between the heterotypic and homotypic interactions^18,17,65^, Strong homotypic interactions may compete with heterotypic interactions, thereby reducing co-partitioning. A comparison of the saturation concentrations (C_sat_) of α-syn and its variants suggests variability in their homotypic condensation, indicating that the variants may exhibit varying homotypic strengths **(Figure 4G)**.

To assess the strength of α-syn’s homotypic interactions directly, we incorporated Acridon-2-ylalanine (Acd), an intrinsically fluorescent unnatural amino acid characterized by long fluorescent lifetimes, high quantum yield, and strong environment sensitivity^66,67,68^, at position 62 of the NAC region in each variant to generate Acd-α-syn or Acd-α-syn^ΔNTD^ or Acd-α−syn^ΔCTD^ **(Figure S5)**. On titrating Acd-α−syn with its WT counterpart, we observed a sharp increase in fluorescence intensity that reached a plateau around 20 μM **(Figure 4H)**. Titration of the other Acd-α-syn variants (CTD or NTD deleted) with their corresponding unlabeled protein fragments also showed similar fluorescence enhancements **(Figure 4I, 4J)** To extract a K_d_ for the homotypic interactions between α-syn, we fit the data to a one-site binding isotherm for all three experiments. Measured dissociation constants were similar in the case of interactions of α-syn and α-syn^ΔCTD^ but were higher for α-syn^ΔNTD^, indicating significantly weaker affinity between two α-syn^ΔNTD^ fragments **(Figure 4K)**. This indicates that NTD deletion reduces homotypic strength, which may promote co-mixing with NP^ΔNTD^ and NP^ΔCTD^ despite the removal of charged blocks.

Since heterotypic interactions appear to be a primary driving force, we used a minimal polymer model from our previous study to understand the relationship between the observed partition coefficient and heterotypic interactions^17^ (see Methods). The model predicts a sigmoidal increase in client partitioning and suggests that partitioning begins above a threshold heterotypic interaction strength (λ_AB_) **(Figure 4L)**. Although the minimal model considers disordered polymer chains, its predictions are consistent with experimental trends, as observed when comparing partition coefficients of NP within the co-condensates formed of α-syn/NP, and different combinations of their variants (as mentioned in Figure 1D, 3D, 3F, 3H) with their respective heterotypic Kd **(Figure 4M)**. This implies that while the minimal model focuses on disordered chains, its underlying principles also apply to multidomain proteins, emphasizing that co-condensation is ultimately governed by a balance of chain-level interactions.

Overall, our findings demonstrate that charge complementarity-driven heterotypic interactions between α-syn and NP primarily drive their co-mixing. Additionally, the modulation of homotypic interactions in α-syn plays a supplementary role in facilitating co-partitioning.

### Accelerated liquid-to-solid transition of droplets in the presence of NP

The spatial organization of protein condensates is a critical aspect that influences the temporal maturation of droplets and subsequent liquid-to-solid phase transition^69^. Previous literature also suggests that transitions between liquid and solid phases can impact the formation of pathological aggregates in other neuronal proteins like TDP-43, FUS and tau^70,71,72^. Therefore, we next investigated the temporal maturation of α-syn and NP separately under homotypic and heterotypic conditions.

We first examined the heterotypic condition by incubating α-syn and NP doped with ∼1% labeled proteins for 24 hours. Additionally, to probe the formation of amyloid-like aggregates inside the droplets during temporal maturation, we added 20 µM Thioflavin T (ThT) to confirm the presence of ThT-positive droplets. ThT is widely used for monitoring amyloid fibril formation; when ThT binds to amyloid fibrils, it produces a robust fluorescent signal. The time-dependent confocal imaging of α-syn and NP (spiked with ThT) in **Figure 5A**, shows that α-syn and NP co-localize from early time points, and the formation of co-condensates is visible at ∼4 hours. In contrast, from 8 hours onward, ThT fluorescence became evident, and all three channels showed co-localization. When the droplets were imaged at ∼12 hours and ∼24 hours, we observed a solid, sea-urchin-like structure with higher ThT partitioning inside the droplets compared with early time points **(Figure 5A,5B)**. These data suggest that condensates formation precedes the accumulation of cross β-structures as the protein mixture forms fibrils. The sigmoidal increase in ThT partition coefficient inside the α-syn/NP droplets is consistent with the formation of amyloid-like aggregates during the liquid-to-solid phase transition **(Figure 5B)**. **Figure 5C** shows a 3D reconstitution generated from confocal image of the co-condensate/aggregate obtained at 24 hours. When a similar experiment was performed with homotypic α-syn droplets incubated with 20 µM ThT, we observed that ThT-positive droplets appeared only after ∼12 hours of incubation **(Figure S6)**. Comparing the measured partition coefficients of ThT over time for homotypic and heterotypic condensates revealed that the accumulation of ThT-positive cross β-structures inside the droplets was much faster in the presence of NP **(Figure 5B)**.

**Figure 5:**
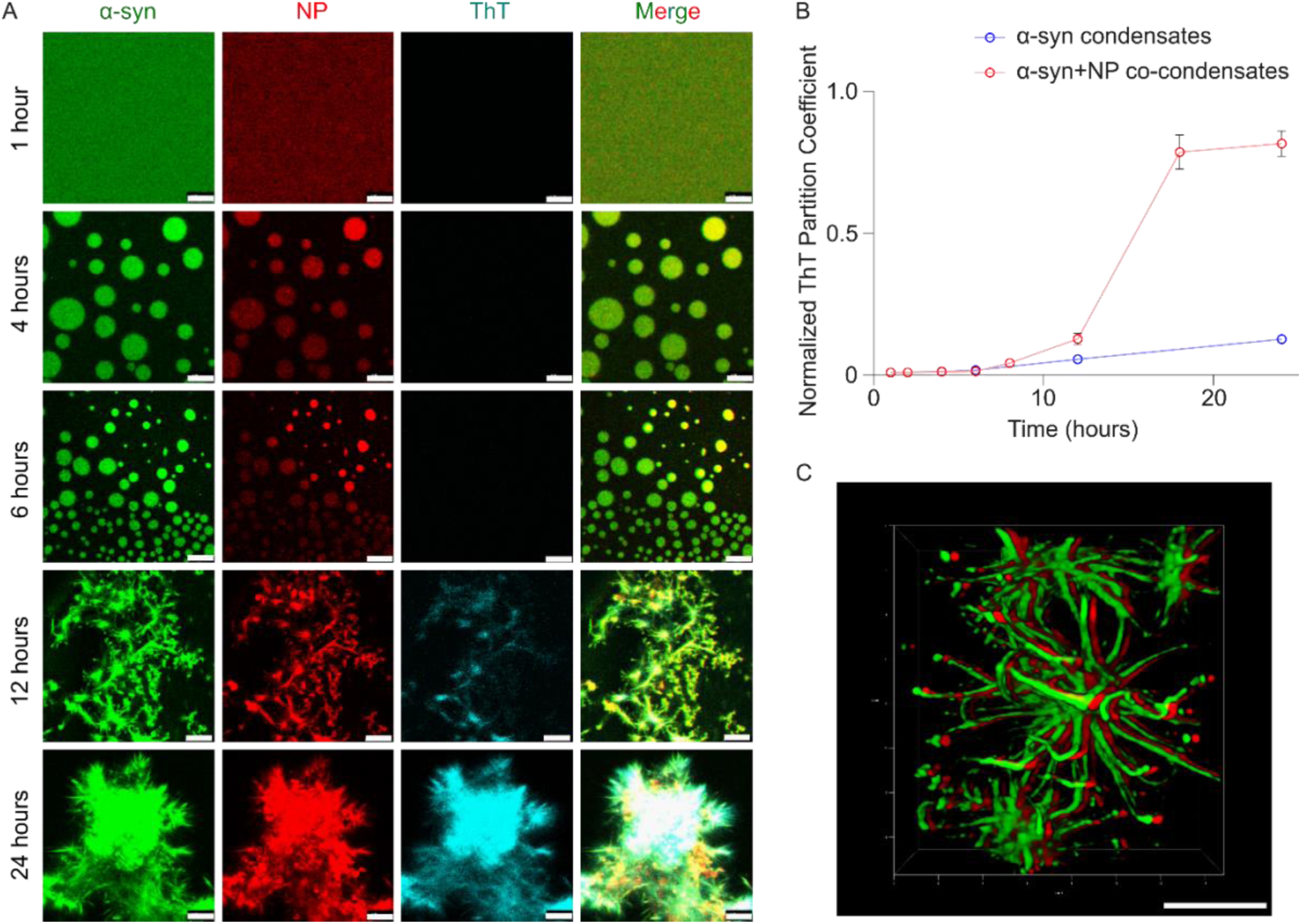
Temporal maturation of the α-syn and NP co-condensates. (A) Time-dependent confocal microscopic images of α-syn (doped with Alexa-488-labeled protein) and NP (doped with Alexa-647-labeled protein) co-incubated with 20μM ThT. The different channels are indicated at the top of the figure panel. The time points of incubation are indicated on the left of the figures. After nearly 12 hours of incubation, ThT-positive solid aggregate-like structures appeared, and at 24 hours of incubation, ThT-positive sea urchin-like structures were observed. Scale bar: 10µm. (B) Plot depicting normalized partition coefficient of ThT in α-syn homotypic droplets and α-syn+NP heterotypic droplets as a function of different incubation time points. Partition coefficients of the α-syn and NP co-condensates are lower at early time points, and an increase in partition coefficient was observed after ∼8 hours. Data are shown as mean± SEM for five independent experiments (n=5). (C) 3D deconvolution of structures obtained after 24 hours of incubation. Scale bar: 5µm.

Using ThT fluorescence, we then studied the in vitro aggregation kinetics of α-syn in the absence and presence of the NP, which confirmed that NP accelerates the fibrillation of α-syn. **(Figure S7A)**. The effect became more pronounced under LLPS conditions, i.e. in presence of crowder and salt. When we examined the fibrils formed after 140 hours by AFM, we found marked differences in the morphology of fibrils between those formed in the absence and presence of NP. The height of the fibrils was in the range of ∼ 3.85nm for α-syn whereas in the presence of NP, it went up to ∼9.23nm **(Figure S7B-C)**.

Overall, our results show that NP accelerates the liquid-to-solid transition of α-syn condensates, leading to faster amyloid-like aggregate formation. This suggests that NP not only promotes phase separation but also influences fibrillation kinetics, which may be relevant to pathological aggregation.

## Conclusions

Abnormal aggregation of amyloid proteins is a hallmark of many neurodegenerative diseases,^73,74^ such as Alzheimer’s and Parkinson’s diseases. Patients with certain neurodegenerative disorders frequently exhibit amyloid pathology associated with another disease, suggesting possible cross-talk between amyloid proteins^75,76,77,78^. Increasing evidence indicates that interactions between different amyloid proteins may contribute to these associations, with heterotypic co-aggregates often exhibiting higher cytotoxicity than homotypic aggregates, leading to greater cellular dysfunction^79,80,81^. While these interactions have been widely studied in the context of amyloid proteins, there is growing interest in exploring how viral infections, particularly SARS-CoV-2, may contribute to neurodegeneration.

In this study, we investigated how the SARS-CoV-2 nucleocapsid protein (NP) influences the phase separation and aggregation of α-synuclein (α-syn), a protein implicated in Parkinson’s disease. α-syn undergoes LLPS, and studies have shown that the nucleation of α-syn aggregation proceeds via metastable bimolecular condensate formation. These liquid droplets eventually transform into pathogenic solid-like aggregates via a liquid-to-solid phase transition, as demonstrated by other amyloidogenic proteins like superoxide dismutase1 (SOD1)^16^, amyloid β^82^, and FUS^82^. Recent studies suggest that, in addition to physicochemical parameters such as temperature and pH^45^, the presence of client molecules (proteins, nucleic acids, or small molecules) can modulate α-syn phase separation and aggregation, influencing the onset and progression of Parkinson’s disease^83,20^.

Our results demonstrate that NP can enhance the phase separation of α-syn in vitro, with co-condensation being depending on both salt valency and ionic strength. We observed a clear dependence on ionic conditions: bivalent ions (e.g., Mg²⁺) promoted stronger phase separation than monovalent ions (e.g., Na⁺), in terms of both droplet number and area of the droplets. Bivalent ions such as Mg^2+^ may also bind α-syn directly, potentially bridging protein chains^84^. Interestingly, elevated magnesium levels found in Parkinson’s patients might contribute to such interactions^85^. Similar salt-sensitive, oppositely charged co-condensation behavior has been observed between ProTα and Histone H1, leading to the formation of highly viscous droplets^31,86^.

Using FCS, we observed that α-syn diffuses ∼4 times faster in homotypic droplets than in heterotypic condensates, suggesting stronger interactions with NP in the dense phase^87^. This restricted dynamics was corroborated by FRAP and fluorescence lifetime data, indicating a more rigid internal environment in the presence of NP.

Electrostatic interactions driving complex coacervation have also been observed between α-syn and other IDR-containing proteins like TDP-43, PrP, and Tau^32,88,89,90,91,92^. These interactions are domain-specific—for example, the N-terminal region of α-syn associates with TDP-43-RNA condensates, while its C-terminal domain (CTD) interacts with Tau and PrP. Our findings similarly indicate that α-syn-NP co-condensation is governed by domain-specific, charge-based interactions. This is supported by molecular simulations and domain deletion studies, highlighting the role of charged regions, in line with patterns seen in transcriptional IDRs^60,93^. Although other non-electrostatic contributions like hydrophobic interactions remain to be investigated, this study lays a foundational framework for understanding the formation of multicomponent condensates.

We also find that in addition to heterotypic interactions, the strength of homotypic interactions within scaffold protein modulates co-condensation. This aligns with prior studies showing that that strong scaffold homotypic interactions can suppress client recruitment, even in the presence of favorable heterotypic interactions^17,18^. Conversely, weakening scaffold homotypic interactions can enhance co-condensation, provided heterotypic interactions exceed a threshold. In agreement, α-syn^ΔNTD^, despite partial absence of charge blocks in domain truncated NP, still co-condensed with NP^ΔNTD^ and NP^ΔCTD^. Homotypic binding assays showed that α-syn^ΔNTD^ had significantly reduced self-interaction, allowing co-condensation via heterotypic contacts. This supports a model in which co-condensation occurs once heterotypic affinity exceeds a critical threshold, and where homotypic strength find-tunes partitioning.

Several studies propose that protein dynamics within liquid droplets are crucial for controlling the maturation process^92,94^. This process is also modulated by the protein’s intrinsic properties and crowding agents^95^. Consistent with our FCS and FRAP data, which revealed slow dynamics of α-syn inside co-condensates, we observed solid-like aggregate formation within ∼ 24 hours, composed of both α-syn and NP. These aggregates can bind the amyloid-specific ThT dye, supporting their fibrillar character. This accelerated liquid-to-solid transition is further evidenced by the sigmoidal increase in ThT partitioning over time.

Overall, our data strongly emphasize the importance of charge complimentary in driving heterotypic condensation and highlight how these interactions within the droplet networks underpin liquid-to-solid transitions **(see Summary Figure 6)**. Based on fluorescence anisotropy and microscopic data, we propose a molecular mechanism in which inter-domain electrostatic interactions facilitate co-condensation. The co-aggregation of α-syn and NP within shared liquid coacervates, as observed here, may underlie their co-localization in disease-related inclusions and could shed light on the mechanistic connection between phase separation and COVID-19 induced Parkinsonism.

**Figure 6:**
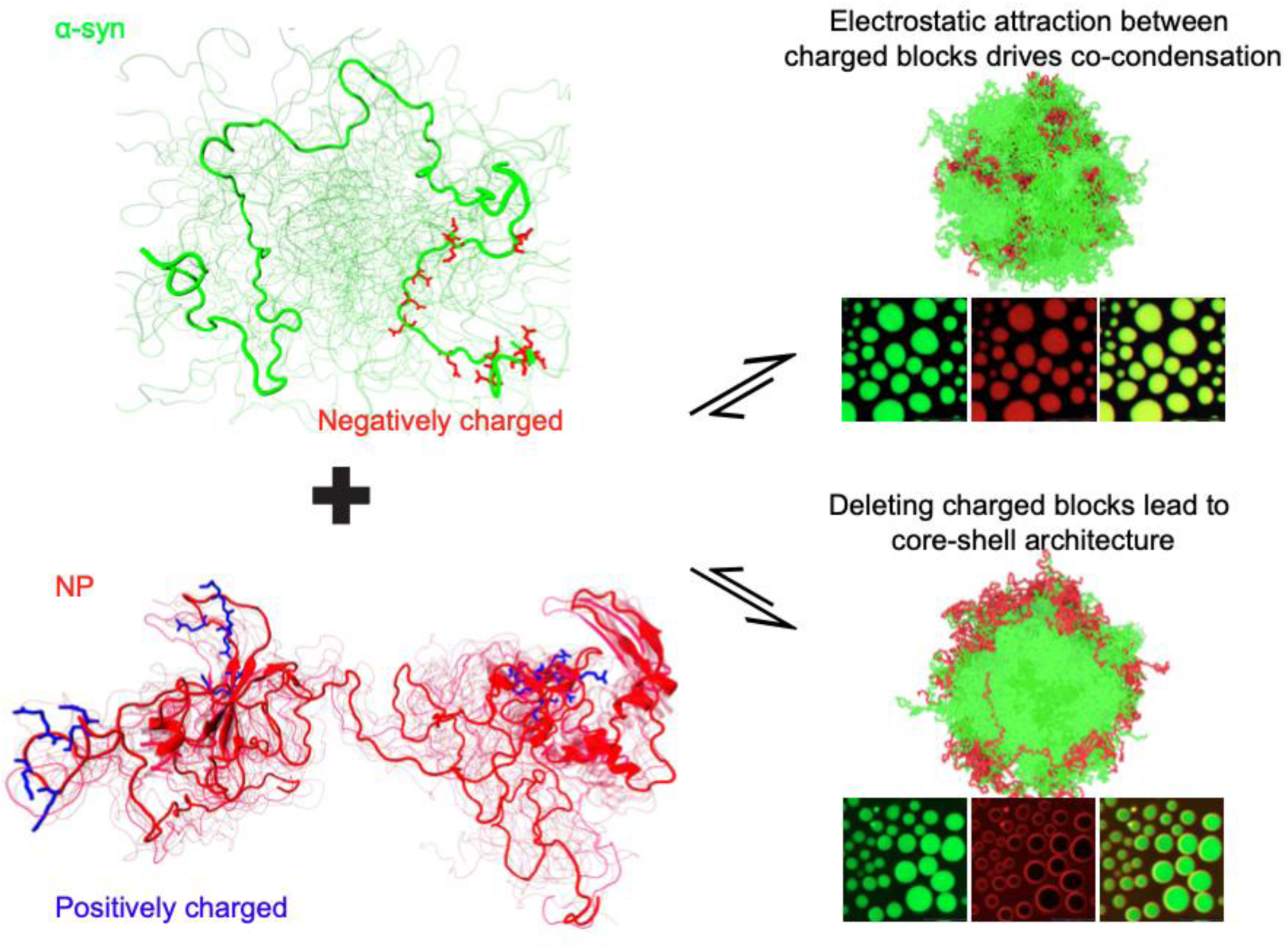
Interactions between oppositely charged blocks drive co-condensation. Heterotypic interactions, driven by electrostatic attraction between oppositely charged blocks, facilitate the co-condensation of α-syn and NP. Removing these charged regions disrupts the interactions, resulting in a core-shell architecture.

## Data Availability

The data presented in this study are available from the corresponding authors upon reasonable request.

## Code Availability

Codes to perform the CG simulations discussed in this work are available at https://glotzerlab.engin.umich.edu/hoomd-blue/. Data analysis and plotting were performed using the MDAnalysis, NumPy, and Matplotlib Python libraries.

## Acknowledgements

The authors acknowledge the CSIR-Indian Institute of Chemical Biology (CSIR-IICB), Kolkata and Academy of Scientific & Innovative Research (AcSIR), India for providing instrumentation facilities. K.C. acknowledges funding from CSIR (OLP-120) for the work done at CSIR-IICB. S.P. thanks Department of Biotechnology (DBT) for awarding fellowships. J.M. gratefully acknowledge the computational resources provided by the Texas A&M High Performance Research Computing (HPRC). K.S. and J.M. thank Busra Ozguney for her valuable feedback on an earlier version of this manuscript. The authors thank Dr. Amy S. Gladfelter (Duke University) for her kind gift of the plasmid pET30b 6his-TEV-SARS-CoV-2-N, which was essential for the initial experimental work.

## Author contributions

**Sumangal Roychowdhury**: Formal analysis; validation; investigation; writing – original draft; figure preparation. **Souradip Paul**: Formal analysis; validation; investigation; writing – original draft; figure preparation. **Kandarp Sojitra**: Formal analysis of simulation data; investigation; methodology; writing – original draft, figure preparation. **Abhishek Mazumder**: Conceptualization; data analysis; writing - original draft. **Sounak Bhattacharjee**: Visualization. **Jeetain Mittal**: Conceptualization; supervision; writing – original draft; project administration. **Krishnananda Chattopadhyay**: Conceptualization; resources; supervision; funding acquisition; writing – original draft.

## Supplemental Information

### Materials

For the purification of full-length α-syn and NP and their variants, LB media, isopropyl-1-thio-β-d-galactopyranoside (IPTG), Tris-HCl, NaCl, imidazole, sodium phosphate monobasic, sodium phosphate dibasic, sodium acetate; from Sigma Aldrich, St. Louis (USA) were used. Ni-NTA resin, protease inhibitor tablet, and SnakeSkin dialysis tube from Thermo Fisher Scientific, Waltham (USA) were used. For the LLPS study, Polyethylene glycol (PEG) 8000 (PEG-8k) was purchased from G-bioscience St. Louis (USA). For protein labelling, the dyes-AlexaFluor488 maleimide and AlexaFluor647 NHS Ester (Succinimidyl Ester), and tris 2-carboxyethyl phosphine (TCEP) were purchased from Thermo Fisher Scientific, Waltham (USA) and Sigma Aldrich, St. Louis (USA) respectively. Apart from these, all other materials used will be mentioned below along with their makers.

### Construct details

For recombinant bacterial expression of human α-syn, the gene encoding for α-syn was cloned in pET21A (+) vector (pET21-ɑ-syn) under the T7 promoter between Nde1 and Xho1 restriction sites. pET21-ɑ-syn G132C mutant was prepared to introduce Cysteine in the ɑ-syn protein for maleimide labelling. pET21-ɑ-synΔNTD (ɑ-syn truncated mutant: Δ1-40) and pET21-ɑ-syn ΔNTD G132C was prepared by deletion mutation on pET21-ɑ-syn and pET21-ɑ-syn G132C plasmids respectively. ɑ-syn ΔCTD construct (ɑ-syn truncated mutant: Δ104-140)-pET21-ɑ-synΔCTD was prepared by deletion mutation on the pET21-ɑ-syn gene, followed by A18C site-directed mutagenesis for maleimide labelling. The codon for Glutamine at 62nd position in pET21-ɑ-syn, pET21-ɑ-synΔNTD, and pET21-ɑ-synΔCTD were mutated to TAG codon to create pET21-ɑ-syn-62TAG, pET21-ɑ-syn-62TAG-ΔNTD, and pET21-ɑ-syn-62TAG-ΔCTD constructs respectively. The primer sets required for synthesizing these constructs are present in Table 1. All the constructs were confirmed by sequencing. The plasmid constructs for purifying recombinant NP (SARS-CoV-2_N_FL) and its domain-deleted variants-NP ΔNTD and NP ΔCTD (SARS-CoV-2_N_deltaNTD and SARS-CoV-2_N_deltaCTD) were a kind gift from Nicolas Fawzi’s lab (Addgene plasmid #157867; Addgene plasmid #157869; Addgene plasmid #157871). pDule2 Mj-Acd-A9 coding for the tRNA_CUA_ and AcdRS was a kind gift from Ryan Mehl’s lab (Addgene plasmid #197652).

**Table 1:**
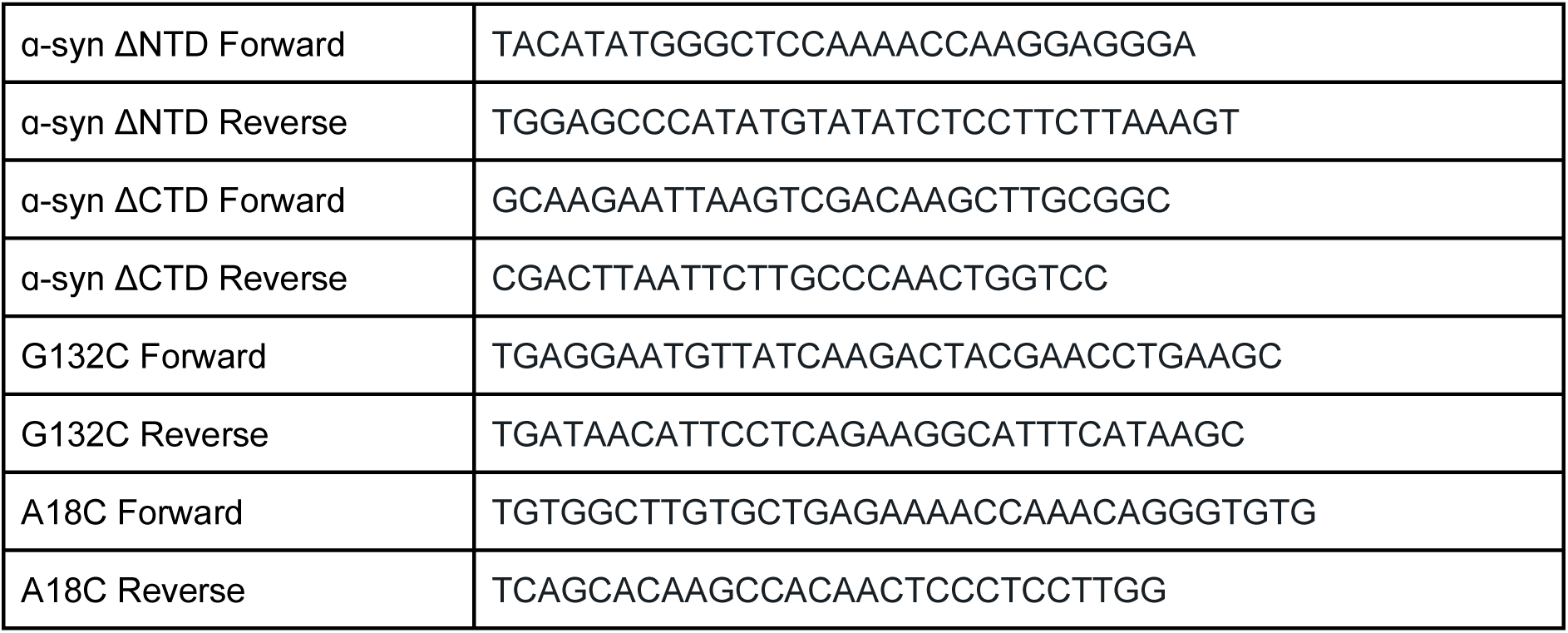
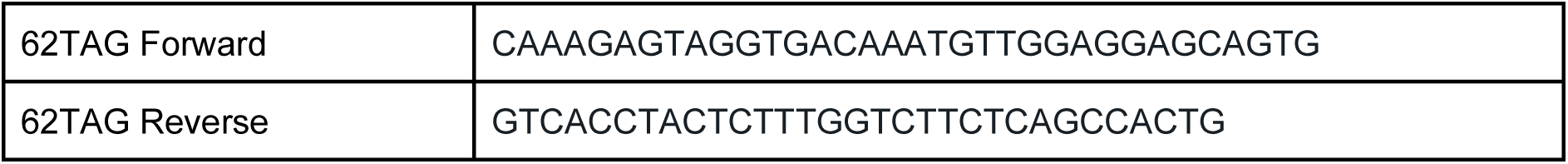
Primers for constructing point mutations and truncations in pET21-ɑ-syn vector.

### α-syn constructs purification and labelling

α-syn and its domain-deleted constructs (α-syn^ΔNTD^ and α-syn^ΔCTD^) were expressed and purified using a previously established protocol^1^. pET21-ɑ-syn constructs were transformed to *Escherichia coli* BL21 (DE3) strain and were then expressed. When the OD_600_ of the culture reached 0.7-0.9, the expression of the protein was induced by adding 1mM isopropyl-β-d-1-thiogalactopyranoside (IPTG) to it, and by incubating the culture at 37^ο^C for 4 hours. Then the cells were harvested using centrifugation and the pellets were resuspended in sonication buffer (20 mM Tris, pH 7.4, 100 mM NaCl and 1 mM PMSF). The resuspended cell pellets were lysed by sonication using short but continuous pulses at 12 Hz for 1 min with a repetition of 14 times. Following sonication, the cell debris was removed from the cell lysates by centrifuging at 14,000 rpm for 45 minutes at 4°C. The lysis suspension was initially brought to 30% saturation with ammonium sulfate and the resulting pellet was discarded. Subsequently, the solution was further saturated to 50% with ammonium sulfate, followed by centrifugation at 20,000 rpm for 1 hour at 4°C. The obtained pellet was resuspended in 20 mM Tris buffer, pH 7.4, and then dialyzed overnight against the same buffer. Following dialysis, the protein sample was filtered using a 30 kDa centrifugal filter. DEAE anion exchange column equilibrated with 20 mM Tris (pH 7.4) was used to purify α-syn from the crude protein. The ion exchange chromatography elution step used a NaCl gradient from 10 mM to 400 mM NaCl. Coomassie-stained SDS-PAGE analyzed the α-syn fractions. It was then concentrated and loaded onto a Sephadex gel filtration column for better purification. The purity of the α-syn protein was 95% using SDS PAGE. Pure α-syn protein was dialyzed against 20 mM sodium phosphate buffer (pH 7.4) buffer for further experiments. The protein concentration was assessed by measuring the absorbance at 277 nm, employing the molar extinction coefficients calculated from the Expasy ProtParam computational tool for each variant, as mentioned in Table 2

**Table 2:** Molar extinction coefficients of α-syn constructs.

|  |  |
| --- | --- |
| α-syn | 5960 M<sup>-1</sup>cm<sup>-1</sup> |
| α-syn<sup>ΔNTD</sup> | 4470 M<sup>-1</sup>cm<sup>-1</sup> |
| α-syn<sup>ΔCTD</sup> | 1490 M<sup>-1</sup>cm<sup>-1</sup> |

### NP constructs purification

NP and its domain-deleted constructs were expressed and purified using already reported protocols with minor modifications^2^. *E. coli* BL21 (DE3) cells harboring MBP-tagged (pTHMT) NP constructs were allowed to grow to an OD_600_ of 0.7 to 0.9, and then 1 mM IPTG was added. The NP variants were expressed at 37^ο^C for 4 hours. The cells were harvested using centrifugation and the pellets were stored at -80^ο^C. The next day, the cells were resuspended in sonication buffer - 20 mM Tris, 1M NaCl, 10mM imidazole pH 8.0. EDTA-free protease inhibitor cocktail tablet was added before lysis via sonication at 12 Hz for 1 min with a repetition of 14 times. Cell debris was removed by centrifugation at 14,000 rpm for 45 min at 4°C. The supernatant was filtered using a 0.2 μm syringe filter and was incubated with Ni-NTA agarose resin at 4°C for 4 hours for binding. The Ni-NTA resin was washed with 20 mM Tris 1 M NaCl 20 mM imidazole pH 8.0. Then, the NP variants were eluted using 20 mM Tris 1 M NaCl 300 mM imidazole pH 8.0. NP eluted fractions were pooled down, buffer exchanged to 20 mM Tris, 1 M NaCl, pH 8.0 and finally concentrated using a 10 kDa centrifugal filter. It was then further purified using size exclusion chromatography by loading it onto a Sephadex gel filtration column. The purity of the fractions was assessed using SDS-PAGE, and fractions which were more than 95% pure were concentrated using a 10 kDa centrifugal filter. These NP variants contained an MBP tag which was further removed for carrying out downstream protocols and experiments. MBP tag was cleaved using TEV Protease (Sigma-Aldrich) as described in the manufacturer’s protocol. In brief, a cleavage reaction containing TEV protease and NP variants at a ratio of 1:100 (w/w) was set up for overnight at 4°C. Then the reaction mixture was loaded onto the Ni-NTA agarose resin column to purify the cleaved NP variants from MBP and hexahistidine-tagged TEV. The NP variants were then further purified using gel filtration. The purity of the NP variants was analyzed using SDS-PAGE and the concentration was measured by recording absorbance at 280 nm. The extinction of each NP variant was calculated using the Expasy ProtParam computational tool as mentioned in Table 3.

**Table 3.** Molar extinction coefficients of NP constructs.

|  |  |
| --- | --- |
| NP | 43890 M<sup>-1</sup>cm<sup>-1</sup> |
| NP<sup>ΔNTD</sup> | 16960 M<sup>-1</sup>cm<sup>-1</sup> |
| NP<sup>ΔCTD</sup> | 26930 M<sup>-1</sup>cm<sup>-1</sup> |

### Acd-ɑ-syn and constructs expression and purification

For experiments in Figures 4H-J and S5, Acd-α-syn, Acd-α-syn^ΔNTD^, and Acd-α-syn^ΔCTD^ were prepared by incorporating the intrinsically fluorescent amino acid Acridon-2yl-alanine (Acd) at position 62 of α-syn or its variants, using unnatural amino acid mutagenesis. Briefly, chemically competent *E. coli* BL21(DE3) cells were co-transformed with plasmids pET21-α-syn-62TAG or pET21-α-syn-62TAG-ΔNTD or pET21-α-syn-62TAG-ΔCTD (constructed from plasmids pET21-α-syn, pET21-α-syn-ΔNTD, or pET21-α-syn-ΔCTD, by use of site-directed mutagenesis to replace codon 62 by an amber codon), and pDule2 Mj-Acd-A9^3^, and spread on a LB agar plate containing 100 μg/mL ampicillin, and 50 μg/mL spectinomycin. A single colony of *E. coli* transformant containing both plasmids was then used to inoculate 10 mL LB broth containing 100 μg/mL ampicillin, and 50 μg/mL spectinomycin and culture were incubated for 16 hours at 37°C with shaking. This culture was added to 1L LB containing 2 mM Acd (TCG Lifesciences Pvt. Limited, India), 100 μg/mL ampicillin, and 50 μg/mL spectinomycin. This 1L culture was incubated in the dark at 37°C with shaking until OD_600_ = 0.7, following which, L-arabinose and IPTG were added to final concentrations of 0.2% and 1 mM, respectively. The culture was further incubated for 4 hours at 37°C with shaking. Cells were harvested by centrifugation (4,000 x g; 20 min at 4°C) and protein was purified following the protocol described for α-syn in the previous section.

### Fluorescent labelling of protein

All α-syn protein variants were labeled using a thiol active fluorescence dye AlexaFluor 488 maleimide (Invitrogen). Proteins containing a single Cysteine residue were dissolved in sodium phosphate buffer pH 8.0 containing 5 mM TCEP and were incubated in ice for 18 hours. Before labeling, the buffer was exchanged to sodium phosphate buffer pH 8.0 containing 0.5 mM TCEP using a 3 kDa Amicon Ultra Centrifugal Filter. The fluorescence dye was dissolved in a minimum amount of DMSO and added to a 1 mg/mL solution of protein under constant stirring. The molar ratio between the protein and dye was 1:10. The reaction mixture was incubated at 20°C for ∼30 min followed by overnight incubation at 4°C. The labelling reaction was then quenched by adding excess β-mercaptoethanol. Excess-free dye and β-mercaptoethanol from the reaction mixture was removed by extensive dialysis in Sodium phosphate (pH 7.4) buffer using SnakeSkin Dialysis Tubing (3 kDa MWCO) followed by column chromatography using a Sephadex G20 column which was pre-equilibrated with 20 mM Sodium phosphate buffer (pH 7.4). All N-protein variants were labeled using an N-terminal fluorescence dye AlexaFluor 647 NHS Ester (Invitrogen, USA) using the manufacturer’s protocol.

### Turbidity assay

α-syn was incubated with 10% PEG-8k, respective salts at different concentrations as mentioned in Figure 1C and S1E. For Figure 1C, a gradual increase in the concentration of NP was added to the reaction mixture containing 10% PEG-8k and 400 mM MgSO_4_. Then turbidity was measured after 4 hours of incubation at 37°C by recording the optical density at 350nm wavelength using a UV spectrophotometer (Thermo Scientific UV-10, USA). In Figure S1E, the turbidity of the mixture containing a fixed ratio of α-syn and NP (20:1) was recorded at different time points. The turbidity data were plotted using GraphPad Prism 8 software.

### Invitro LLPS assay

To confirm LLPS, imaging of droplets was done using the confocal microscope. The proteins were doped with 1% respective labeled protein in a ratio of 99:1, and the LLPS was facilitated using different concentrations (w/v) of PEG-8k and different salts of concentrations for optimization of standard conditions as mentioned in Figures 1 and S1. After initial screening final conditions selected are α-syn and NP constructs (20:1), 10% PEG-8k and 400 mM MgSO_4_. These conditions were used for all further experiments involving different constructs of α-syn and NP. The solutions were incubated for 4 hours at 37 °C with constant shaking at 180 rpm. Then the sample was drop cast on a 22 mm grease-free extensively cleaned coverglass (Blue-Star, India). Then imaging was performed using an inverted confocal microscope (Leica TCS SP8, Germany) equipped with a 63X (1.4 NA) oil immersion objective lens. AlexaFluor 488 maleimide tagged α-syn was excited using an Argon 488nm line laser and AlexaFluor 647 NHS Ester tagged NP was excited using the HeNe 633nm line laser. After image acquisition at a frame size resolution of 1024 X 1024 pixels, the images were processed and analyzed using Image J software (Developer-National Institute of Health, USA) to find the intensity of fluorescence signal in the dense and dilute phases, size and number of droplets.

### Fluorescence Recovery After Photobleaching

The ɑ-syn homotypic and the ɑ-syn/NP heterotypic droplets were imaged using the protocol above. FRAP experiments were performed to understand the material properties of the droplets. Here, a specific region within the droplet was completely bleached using a defined ROI for 15 seconds by 80% laser power of an Argon 488 nm laser line. The ROI was calculated as 10% of the total droplet area. Then recovery was observed for 130 minutes with an interval of 2 seconds. For the FRAP experiment on NP in the co-condensate, 80% laser power of HeNe 633 nm laser was used; and parameters were pre-bleach time of 2 seconds, bleach time of 6.48 seconds, and recovery time of 91 seconds with an interval of 1 second. Post-bleaching, the average fluorescence intensity in defined ROIs was plotted against respective recovery time using GraphPad Prism 8 software to determine the percentage recovery compared to the pre-bleached fluorescence intensity.

### Confocal imaging of Merocyanine 540 (MC540) localization in condensates

Merocyanine 540 was used as a fluorogen to probe the relative hydrophobicity of the homotypic ɑ-syn droplets and heterotypic α-syn+NP co-condensates. The homotypic and heterotypic LLPS was facilitated using the aforementioned standardized protocol. 50μM MC540 was added to the reaction mixture and no labeled proteins were used as it may lead to crosstalk of the fluorescence signal of MC540. Then confocal microscopy imaging was performed using an inverted confocal microscope (Leica TCS SP8, Germany) and a 532 nm laser line was used to excite MC540 internalized droplets. All other imaging parameters were the same as the imaging protocol of the Invitro LLPS assay mentioned earlier. The partitioning of MC540 was determined by calculating the ratio of fluorescence intensity signal of MC540 inside and outside the condensates, measured using Image J software (Developer-National Institute of Health, USA) and plotted using GraphPad Prism 8 software.

### Atomic Force Microscopy (AFM)

Aggregating samples of α-syn and co-incubated samples of α-syn + NP were aliquoted after prolonged incubation for ∼140 hours at 37°C and diluted 20 times with Milli-Q water. 7 μL diluted sample was drop cast on a freshly cleaved mica sheet (muscovite mica, Biolyst/Electron Microscopic Science, Pennsylvania, USA). The aggregates on the mica sheet were carefully rinsed twice with MilliQ water and then dried using a stream of nitrogen. AFM images were acquired at room temperature using a Asylum Research AFM (Oxford instruments, UK) with silicon probes. The standard tapping mode was used to image the morphology of aggregates. The nominal spring constant of the cantilever was kept at 20–80 N/m. The spring constant was calibrated by a thermal tuning method. A standard scan rate of 0.5 Hz with 512 samples per line 6 was used for imaging the samples. A single third-order flattening of height images with a low pass filter was done followed by section analysis to determine the dimensions of aggregates using previously described methods^4^.

### Thioflavin T (ThT) assay

Aggregation kinetics was determined for α-syn and both α-syn+NP (in a 20:5 standardized ratio). The concentrations used for the aggregation assay were the same as the in vitro LLPS assay-α-syn (100 μM) or α-syn and NP (100 μM + 5 μM) present in 20 mM sodium phosphate buffer (pH 7.4). The reactions for the aggregation kinetics assay were set both in normal condition and LLPS condition (containing 10% PEG-8k and 400 mM MgSO_4_) as mentioned in Figure S7A. Aliquots of the reaction mixture were withdrawn at each time point in the aggregation pathways and were diluted 10 times in 20 mM Sodium phosphate buffer (pH 7.4) and then, ThT was added to the diluted reaction mixture in a 1:3 molar ratio (α-syn: ThT) to reach a total volume of 500 μL. Steady-state fluorescence measurements were recorded in a quartz cuvette (Hellma, Sigma-Aldrich, USA) of path length 1 cm using a Photon Technology International (PTI) fluorescence spectrometer (Horiba, Japan) with an excitation wavelength of 450 nm, emission wavelength of 485 nm and an integration time of 0.1 s averaging over 3 times. Three independent experiments were performed for each set (N=3). Then ThT fluorescence intensities of each experiment were plotted against their respective time points using OriginPro2024 software.

### Steady-state fluorescence anisotropy

Steady-state fluorescence anisotropy experiments were performed to measure the strength of heterotypic interaction between α-syn and NP constructs. 100 nM of AlexaFluor 488 labeled ɑ-syn was titrated with increasing concentration of NP in the reaction mixture containing 20 mM sodium phosphate buffer pH 7.4, and was incubated at 25°C for 30 minutes to ensure maximum binding. Quartz cuvette of path length 1cm (Hellma, Sigma-Aldrich, USA) was treated with 1mg/mL BSA in 20mM sodium phosphate buffer pH 7.4 to ensure protein samples do not get stuck to the quartz surface. Then anisotropy measurements of the ɑ-syn were performed at 25 τC in a quartz cuvette using a PerkinElmer LS 55 Luminescence spectrometer (USA). The excitation wavelength was set to 490 nm and the emission signal was recorded at emission maxima of 525 nm with an integration time of 10 seconds. Three independent experiments were performed for each heterotypic strength assay involving different α-syn and NP constructs as mentioned in Figure 4A-C (N=3). The fluorescence intensities were measured in all four excitation and emission polarizer combinations-Ex_vertical_-Em_vertical_, Ex_vertical_-Em_horizontal_, Ex_horizontal_-Em_vertical_, and Ex_horizontal_-Em_horizontal_. Then the anisotropy value (r) was calculated as follows:

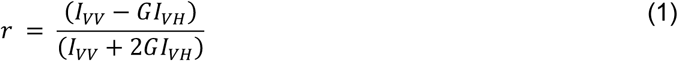

where I_VV_, I_VH_, I_HV_, and I_HH_ are fluorescence intensity for vertical excitation and vertical emission, vertical excitation and horizontal emission, horizontal excitation and vertical emission, and horizontal excitation and horizontal emission respectively. The Grating factor (G) is defined as

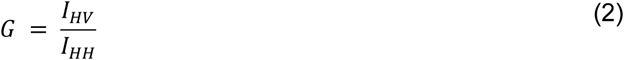

The anisotropy values were plotted against the respective titration concentration using GraphPad Prism 8 software and the binding isotherm was fitted using the following equation:

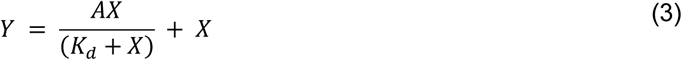

where X is the concentration of titrated NP constructs, Y is the anisotropy values of ɑ-syn at different titrations, A is the maximum binding measured by the maximum anisotropy value, and K_d_ is the equilibrium dissociation constant.

### Fluorescence equilibrium stoichiometry titration

To quantify the homotypic interaction, we exploited genetic code expansion methodology to label ɑ-syn. ɑ-syn does not have intrinsic Trp, so, a fluorescent unnatural amino acid Acridon-2-ylalanine (Acd) was inserted at the 62^nd^ position using the earlier mentioned protocol. In brief, the codon for glutamine at the 62^nd^ position of ɑ-syn was mutated to amber (UAG), a minor stop codon in *E.coli*, which also happens to code for Acd. This position of substitution (Q62Acd) is present in the NAC region and, hence available for all three ɑ-syn variants. Then 200 nM Acd-ɑ-syn constructs were individually titrated with respective unlabeled ɑ-syn construct in 20 mM Sodium Phosphate Buffer pH 7.4. The reaction mixture was incubated at 25°C for 30 minutes to optimize binding efficiency. The fluorescence intensity was measured in a quartz cuvette (Hellma, Sigma-Aldrich, USA) of path length 1 cm, using a Photon Technology International (PTI) fluorescence spectrometer (Horiba, Japan). Three individual sets of experiments were performed for each homotypic strength assay as mentioned in Figure 4H-J (N=3). The excitation wavelength for steady-state fluorescence measurements was 370 nm and the emission was recorded at 421 nm with an integration time of 1 second. Upon titrating unlabeled ɑ-syn against a fixed concentration of Acd-ɑsyn, the fluorescence intensity gradually increased and became saturated at higher concentrations. The fluorescence intensity was plotted against the respective titration concentration in OriginPro2024 software to generate a binding isotherm which was then fitted using the following equation:

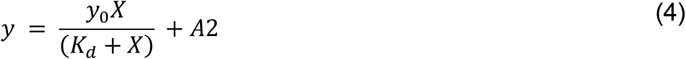

where X is the concentration of titrated ɑ-syn constructs, y is the normalized fluorescence intensity values of ɑ-syn at different titrations, *y*_0_ is maximum intensity measured at x=0, K_d_ is the equilibrium dissociation constant and A2 is any non-specific background.

### Lifetime of protein condensates

50 nM labeled ɑ-syn along with 100 µM unlabeled ɑ-syn or 100 µM unlabeled ɑ-syn+ 5 µM unlabeled NP were subjected to LLPS conditions and droplets were visualized under a confocal microscope Nikon Ti2 for Time-Domain FLIM experiments. The time-resolved fluorescence measurements were made using Alba’s time-correlated single photon counting (TCSPC) setup (ISS Inc., Champaign, Illinois). Measurement samples were excited using a 488 nm QuixX picosecond pulsed laser made by Omicorn-Laserage Laserprodukte GmbH. The repetition rate of the laser was set to be 20 MHz. The laser was linearly polarized in the vertical direction, and a linear polarization cleanup filter (DPM-100-VIS by Meadowlark Optics) was used to improve the extinction ratio further. For FLIM measurements, the fluorescence emission was detected by a single photon avalanche diode (SPAD) detector (SPD-100-CTC by Micro Photon Devices) after the 530/43-nm band-pass filter (Semrock). The FLIM data were analyzed using the ISS 64-bit VistaVision software. In the FLIM data analysis, the software allows the single or multiexponential curve fittings on a pixel-by-pixel basis using a weighted least-squares numerical approach. The single-exponential model was used for fitting the lifetime data of ɑ-syn and ɑ-syn+NP condensed phase with the instrument response function, estimated by taking the first derivative of the rising of the decay. The histograms of the lifetime distribution were plotted using OriginPro2024 software.

### Fluorescence Correlation Spectroscopy (FCS)

The translational dynamics of the droplets were measured using FCS. FCS analyzes fluctuations in fluorescence intensity within the confocal volume to determine the diffusion times of fluorescent molecules present in nanomolar concentration. ɑ-syn homotypic condensates were prepared using the previous protocol where 100 µM unlabeled ɑ-syn was doped with 15 nM Alexa-488 labeled ɑ-syn in the presence of 10% PEG-8k and 400 mM MgSO_4_. Similarly, heterotypic condensates were also prepared but only labeled ɑ-syn was added as a fluorescent probe. The FCS measurement was performed using an ISS Alba FFS/FLIM confocal system (Champaign, IL, USA), integrated with a Nikon Ti2U microscope equipped with the Nikon CFI PlanApo 60X/ 1.2NA water immersion objective. The LLPS sample was drop cast on the 22 mm grease-free coverglass (Blue-Star, India) and was allowed to settle for ∼10 minutes so that the droplets formed will get immobilized on the coverslip. Then, droplets were imaged at the frame size of 512 X 512 pixels and the pixel dwell time was set to 0.1 ms. ROIs were set on the droplets which were larger than the focal spot and the Z-axis was tuned to measure the diffusion inside the droplet and not on the surface. The ROIs were excited one at a time using a 488-nm picosecond pulsed diode laser and FCS measurements were acquired. The fluorescence emission was detected using a SPAD (Single Photon Avalanche Detector) detector and the 530/43-nm band-pass filter. The FCS correlation curves obtained were fit to the 3D Gaussian 1-component diffusion model using VistaVision software (ISS Inc., USA). The fitted FCS correlation curves were plotted using GraphPad Prism 8 software. The beam waists in the radial and axial dimensions were calibrated using a standard fluorescence dye, Rhodamine 6G (R6G), in water with a known diffusion coefficient of 2.8 × 10⁻⁶ cm²/s. The diffusion coefficients (D) of the dense and dilute phases of homotypic and heterotypic droplets were determined by measuring the diffusion time (τ_D_) and applying the relation from^5^:

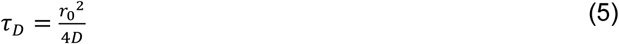

Where, *r*_0_ = Where r0 is the lateral radius of the hypothetical ellipsoid observed volume.

The FCS data collected were normalized to free dye using a method previously described by Chattopadhyay et al.^6^. In the 3D Gaussian diffusion model, which considers a single type of diffusing molecule (the 3D Gaussian 1-component model, excluding triplet state contributions), the correlation function G(τ) is defined by the following equation:

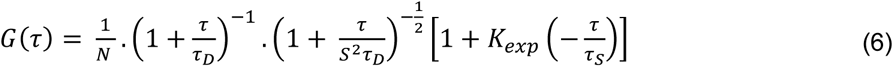

where *τ*_*D*_ represents the diffusion time of the molecules, N is the average number of molecules within the observation volume, and S is the structural parameter that defines the ratio between the radius and the height of the confocal volume. The value of *τ*_*D*_, obtained by fitting the correlation function, is related to the diffusion coefficient (D) of a molecule through the following equation:

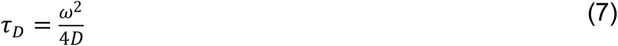

where ω is the size of the observation volume.

### Time-dependent maturation study

The time-dependent maturation experiment was done with both α-syn homotypic droplets and α-syn - NP heterotypic droplets for ∼24 hours. For homotypic droplets, 100 µM unlabeled α-syn samples were spiked with 1% Alexa 488-labeled α-syn and 20 µM ThT dye. The protein sample was incubated with 10% PEG-8k, 400mM MgSO_4_ at 37°C with constant shaking at 180 rpm for 24 hours. Samples were taken out at indicated time points and imaged in a confocal microscope (Leica SP8, Germany). α-syn droplets were imaged using Argon 488 nm laser line and for the imaging of ThT fluorescence, we used a 458 nm laser line. Similarly, for heterotypic droplets, 100 µM unlabeled α-syn (spiked with 1% Alexa 488-labeled α-syn), 5 µM unlabeled NP (spiked with 1% Alexa 647-labeled NP) and 20 µM ThT dye was incubated with 10% PEG-8k, 400 mM MgSO_4_ at 37°C with constant shaking at 180 rpm for 24 hours. Samples were taken out at indicated time points and imaged in a confocal microscope (Leica SP8, Germany). For imaging NP we used a HeNe 633 nm laser line. All the fluorescence images were taken in 63X oil immersion objective (NA - 1.4) with a resolution of 1024 x 1024 and analyzed using ImageJ (NIH, USA). Five individual sets of experiments were performed for the time-dependent maturation study (N=5). The ThT partition coefficient at different time points was calculated from the ratio of the intensities in the dense (inside) and dilute (outside) phases of the condensates. ThT partition coefficients were plotted against the respective time points using GraphPad Prism 8 software. The 3D reconstitution of the heterotypic droplet/aggregate image at 24 hours was done using Huygens Deconvolution software (Scientific Volume Imaging B.V.).

### Coarse grained simulations

Coarse-grained (CG) coexistence simulations were carried out using the HOOMD-Blue 2.9.3 software package^7^, following the protocol of a previously established CG simulation approach for proteins^8,9^.

Protein chains were modelled using the HPS-Urry model^10^. The bonded interactions between amino acids were described by a harmonic potential:

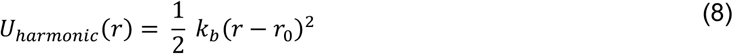

where k_b_ is the harmonic spring constant (20 kcal/(mol Å^2^)), and r_0_ represents the equilibrium bond length (3.8 Å). Nonbonded interactions between monomers were governed by a modified Lennard–Jones (LJ) potential^11^:

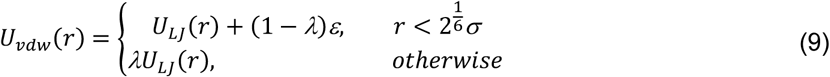

where λ modulates the attractive well depth, and U_LJ_(r) follows the standard Lennard–Jones form:

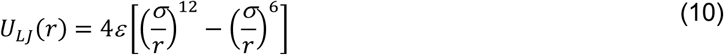

The interaction parameter (ε) was determined individually for each protein to ensure that the simulated dilute-phase concentration aligned with experimental saturation concentrations. For α-syn, ε was set to 0.267 kcal/mol, while for NP, it was assigned a value of 0.194 kcal/mol. The cross-term interaction parameter was derived based on geometric mean. To model NP, the disordered regions (N_IDR_, linker, and C_IDR_) were kept flexible, whereas the folded domains (NTD and CTD) were constrained using the hoomd.md.constrain.rigid function^12^. In the case of α-syn, all regions were modeled as flexible.

The initial configuration was set up with 405 α-syn chains and 20 NP chains in a cubic simulation box measuring 15 nm × 15 nm × 15 nm, maintaining an experimental molar ratio of α-syn:NP = 20:1. Simulations were conducted at 300 K with a time step of 10 fs. A Langevin thermostat was employed, where the residue friction coefficient was defined as γ=m_i_/t_damp_, with m_i_ representing the mass of each residue and t_damp_ set to 1000 ps. The system was simulated in an NVT ensemble for 5 μs, with the first 1 μs discarded from the analysis as equilibration. The pairwise residue index contact map was constructed by defining a contact between two residues, *i* and *j*, if their distance was less than 1.5σ. The contact propensity, n_contact_, was calculated as

<n_ij_>, where n_ij_=1 if the distance criterion was met and n_ij_=0 otherwise. In the contact maps, the contact propensities were represented as -ln(n_contact_ /n_max_), where n_max_ denotes the highest observed contact propensity for any residue pair. To identify charged blocks, we computed the net charge within a sliding window of 10 residues^13^. A window was designated as positively or negatively charged if its net charge was ≥ +5 or ≤ -5, respectively. Adjacent windows meeting this criterion were merged to define continuous positively or negatively charged blocks. In Figure 6, the structural ensemble of α-syn was obtained from PED00024, while the NP structures were generated using MODELLER^14^. The surface electrostatic potential was computed using Adaptive Poisson-Boltzmann Solver (APBS) software^15^. All structural snapshots were generated using VMD 1.9.3^16^ and UCSF ChimeraX^17^.

To compare heterotypic interaction strength with partition coefficients (Figure 4L), we used data from our previously developed minimal polymer model^18^. Briefly, the system consisted of two polymer chains, Scaffold A and Client B, each 20 monomers in length. Bonded interactions between monomers were modelled using a harmonic potential, while nonbonded interactions were described by a modified LJ potential. In this model, the σ was set to 5 Å, and the ε was fixed at 0.20 kcal mol⁻¹. The scaffold’s homotypic strength (λ_AA_) was held constant at 1, whereas the client’s homotypic strength (λ_BB_) and heterotypic interaction between the scaffold and client (λ_AB_) were systematically varied from 0 to 1. To explore the parameter space, we performed scanning simulations by keeping λ_AB_ constant while varying λ_BB_ from 0 to 1 in increments of 0.02, and vice versa. Additional scans were conducted along linear lines above and below the diagonal to examine different λ_BB_–λ_AB_ combinations. Each simulation was run for 100 ns. Partitioning was quantified by computing the partition coefficient as the ratio of B’s concentration inside A to its concentration in the surrounding solution, rather than the dilute phase, to avoid numerical issues at low values.

---

**Figure S1:**
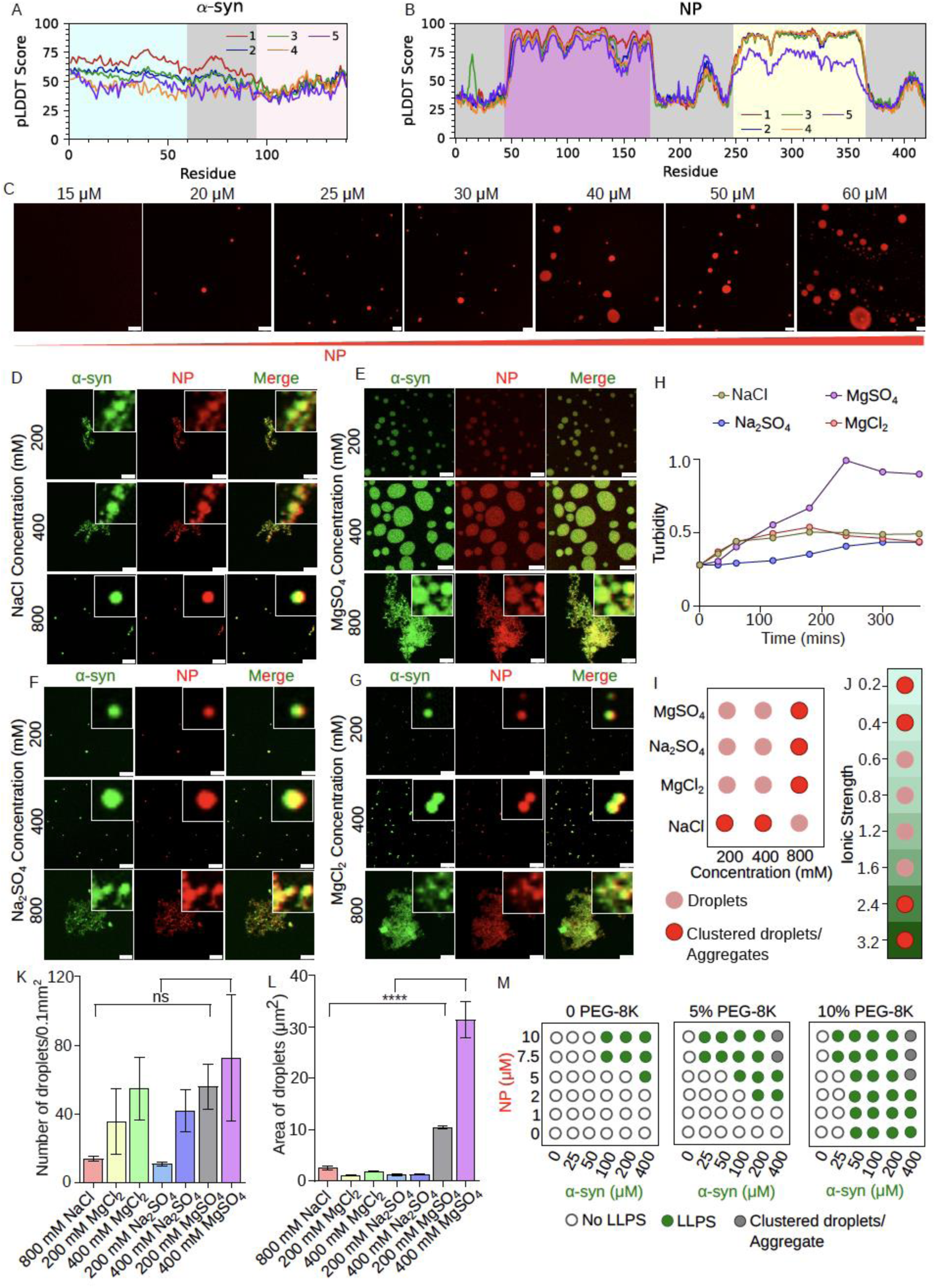
Influence of ionic strength and crowding on co-condensation. (A) pLDDT score of top 5 models predicted using ColabFold^19^ of (A) ɑ-syn and (B) NP (C) Confocal microscopic images of Alexa-647 labeled doped NP at different unlabeled protein concentrations for the determination of saturation concentration (C_sat_). The observed C_sat_ was 20µM. Scale bar: 10µm. Three independent experiments were done (n=3). (D) Confocal microscopic images of co-condensates of Alexa-488 labeled ɑ-syn and Alexa-647 labeled NP (Molar ratio 20:1) in the presence of 10% PEG-8k and NaCl after ∼ 4h of incubation at 37τC and 180 rpm. Scale bar: 10µm. The salt concentrations are indicated on the left of each panel. Three independent experiments were done (n=3). (E) Confocal microscopic images of co-condensates of Alexa-488 labeled ɑ-syn and Alexa-647 labeled NP (Molar ratio 20:1) in the presence of 10% PEG-8k and MgSO_4_ after ∼ 4h of incubation at 37τC and 180 rpm. Scale bar: 10µm. The salt concentrations are indicated on the left of each panel. Three independent experiments were done (n=3). (F) Confocal microscopic images of co-condensates of Alexa-488 labeled ɑ-syn and Alexa-647 labeled NP (Molar ratio 20:1) in the presence of 10% PEG-8k and Na_2_SO_4_ after ∼ 4h of incubation at 37τC and 180 rpm. Scale bar: 10µm. The salt concentrations are indicated on the left of each panel. Three independent experiments were done (n=3). (G) Confocal microscopic images of co-condensates of Alexa-488 labeled ɑ-syn and Alexa-647 labeled NP (Molar ratio 20:1) in the presence of 10% PEG-8k and MgCl_2_ after ∼ 4h of incubation at 37τC and 180 rpm. Scale bar: 10µm. The salt concentrations are indicated on the left of each panel. Three independent experiments were done (n=3). (H) Plot of time-based turbidity measurement in the presence of different salts at 400mM concentration. Data is shown as mean± SEM. Three independent measurements were done with similar observations (n=3). MgSO4 showed the highest turbidity after ∼ 4h of incubation. (I) Schematic representation of droplet and clustered droplet/aggregate formation at different salt concentrations for different types of salts as observed from confocal microscopic experiments (Figure D-G). The pink circle indicates droplets, and the red circle indicates clustered droplets/aggregate. (J) Range of ionic strength at which droplet formation is mostly favored. The intermediate ionic strength ranges from 0.6-1.6 favors the droplet formation. (K) Plot depicting the number of droplets observed in confocal microscopic images for heterotypic co-condensates of ɑ-syn and NP at indicated salt concentrations of different types of salts. Data is shown as mean± SEM (n=3). “ns” indicates non-significant. P values were determined by non-parametric t-test. (L) Plot depicting the area of droplets observed in confocal microscopic images for heterotypic co-condensates of ɑ-syn and NP at indicated salt concentrations of different types of salts. Data is shown as mean± SEM (n=3). **** indicates P value < 0.0001. Statistical significance was established by non-parametric t-test. (M) State diagram indicating the effect of crowding agent, PEG-8k on co-condensation of ɑ-syn and NP. The added percentage of PEG-8k is indicated at the top of the state diagram. The hollow circle indicates no phase separation, the solid green circle indicates complete co-localization via phase separation and the solid grey circle indicates co-aggregates. Three independent experiments were done (n=3).

**Figure S2:**
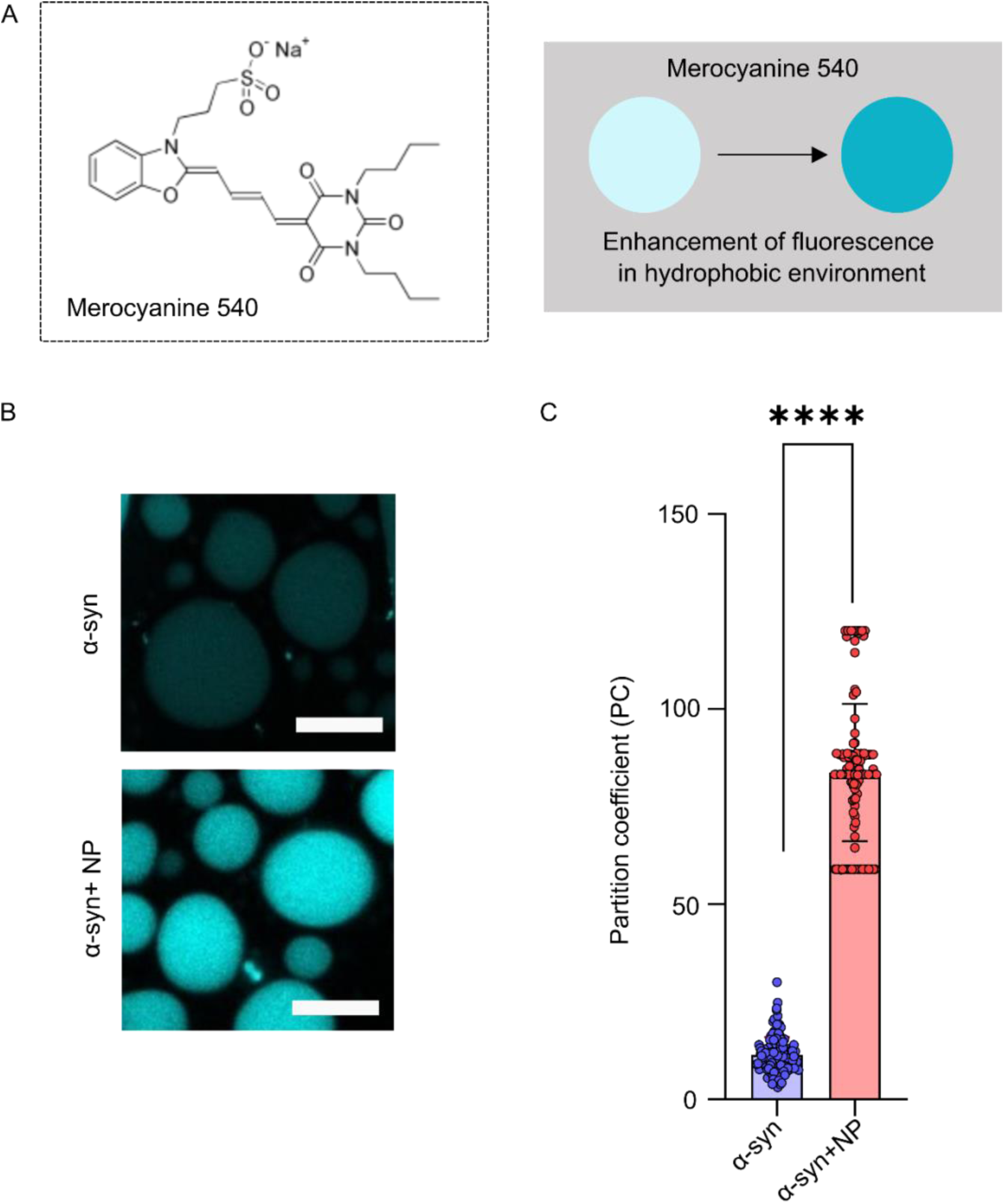
Probing the droplet environment by Merocyanine540. (A) Structure of Merocyanine540 and schematic representation of environment sensitivity. Schematic representation showing the mode of action of MC540. Enhancement of fluorescence of the dye is observed for a more hydrophobic environment. (B) Confocal microscopic images of condensates of unlabeled ɑ-syn (100 µM), co-condensates of unlabeled ɑ-syn and unlabeled NP in the presence of 10% PEG-8k, 400 mM MgSO_4_ ∼ 4h of incubation at 37τC and 180 rpm and spiked with MC540. Scale bar: 10µm. Three independent experiments were done (n=3). (C) Partition coefficient of MC540 for homotypic condensates of ɑ-syn and heterotypic condensates of ɑ-syn and NP. The partition coefficient was higher for full-length heterotypic condensates of ɑ-syn and NP, which indicates a more hydrophobic environment compared to homotypic ɑ-syn droplets. Three independent experiments were done (n=3). For partition coefficient calculation n=141 droplets were measured for homotypic condensates of ɑ-syn and n=135 droplets were measured for heterotypic condensates of ɑ-syn and NP. Data is shown as mean± SD. **** indicates P value < 0.0001. Statistical significance was established by a non-parametric t-test.

**Figure S3:**
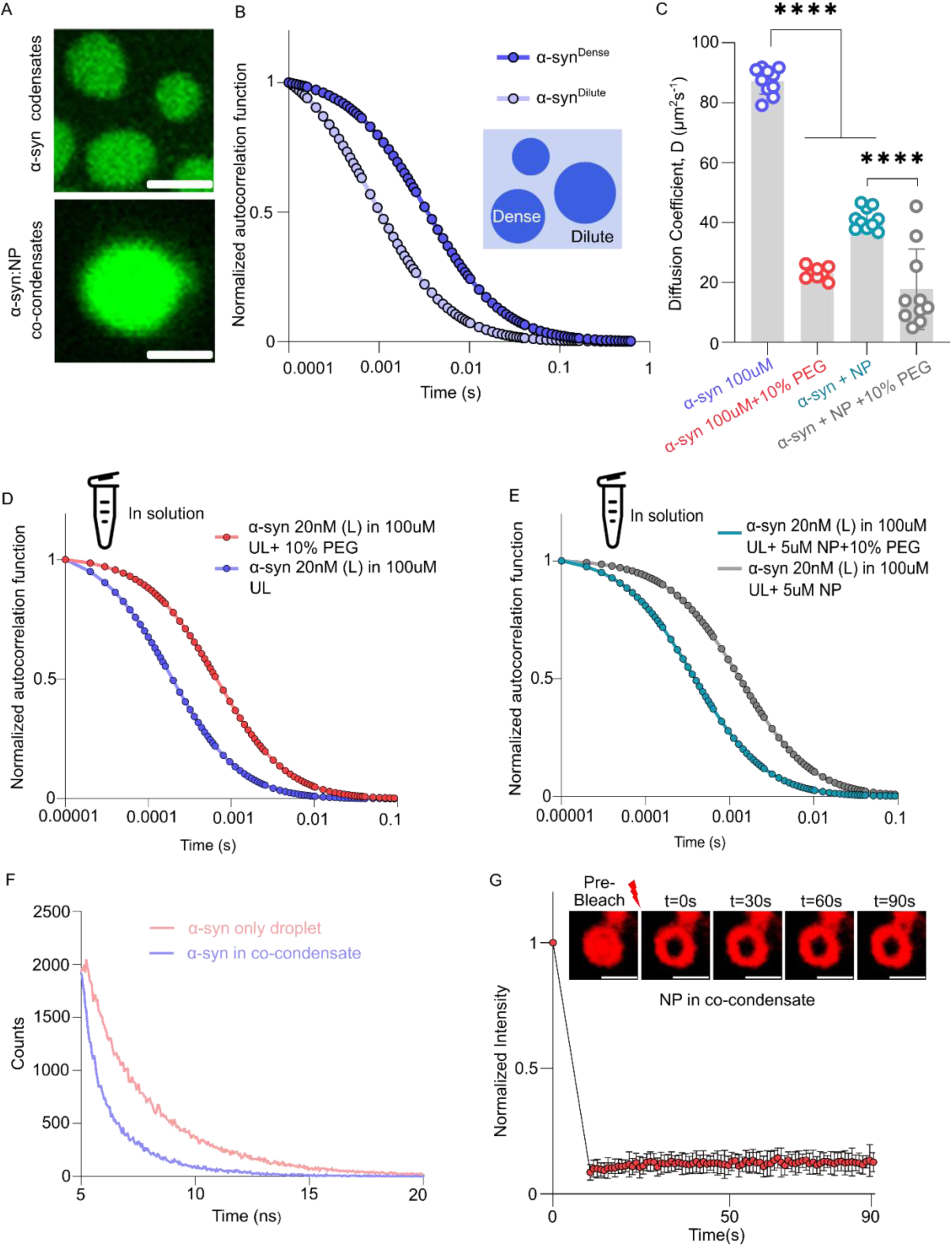
Probing dynamics of ɑ-syn inside the condensate. (A) Representative confocal microscopic images taken in FCS setup of Alexa-488 labeled ɑ-syn homotypic condensates and heterotypic co-condensates of ɑ-syn and NP doped with only Alexa-488 labeled ɑ-syn in the presence of 10% PEG-8k and 400 mM MgSO_4_ after ∼ 4h of incubation at 37τC and 180 rpm. Scale bar: 10µm. Three independent experiments were done (n=3). (B) Amplitude normalized autocorrelation function obtained from FCS of ɑ-syn in the homotypic condensates. Intense color indicates the dense phase of the condensates and light variant indicates the dilute phase of the condensates as indicated in the schematic representation shown in the inset. The shift in the autocorrelation function indicates the increase in the diffusion time as observed in the dense phase compared to the dilute phase. (C) Plot depicting a comparison of diffusion coefficients obtained from FCS measurements of ɑ-syn only and in addition with NP in solution phase, with or without 10% PEG-8k. Data is shown as mean± SD (n=10). **** indicates P value < 0.0001. Statistical significance was established by a non-parametric t-test. (D) Amplitude normalized autocorrelation function obtained from FCS of ɑ-syn in the solution phase to show the effect of crowding on the diffusion coefficient of ɑ-syn. “L” represents labeled ɑ-syn and “UL” represents unlabeled ɑ-syn. (E) Amplitude normalized autocorrelation function obtained from FCS of ɑ-syn with addition of NP in the solution phase to show the effect of crowding on the diffusion coefficient of ɑ-syn. “L” represents labeled ɑ-syn and “UL” represents unlabeled ɑ-syn. (F) TSCPC decay curve obtained from TD-FLIM imaging for ɑ-syn in homotypic and heterotypic co-condensates. The pink color indicates ɑ-syn in homotypic condensates and the violet color indicates ɑ-syn in heterotypic condensates. (G) Inset: A representative droplet image at indicated time points of FRAP measurements for NP in heterotypic droplets of α-syn and NP. Scale bar: 2µm FRAP kinetics curve for NP in heterotypic droplets of α-syn and NP. The mobile fractions for NP in heterotypic droplets of α-syn and NP was ∼ 5%, indicating almost immobility of NP in the co-condensates. Data is shown as mean± SD (n=3) for three independent experiments.

**Figure S4:**
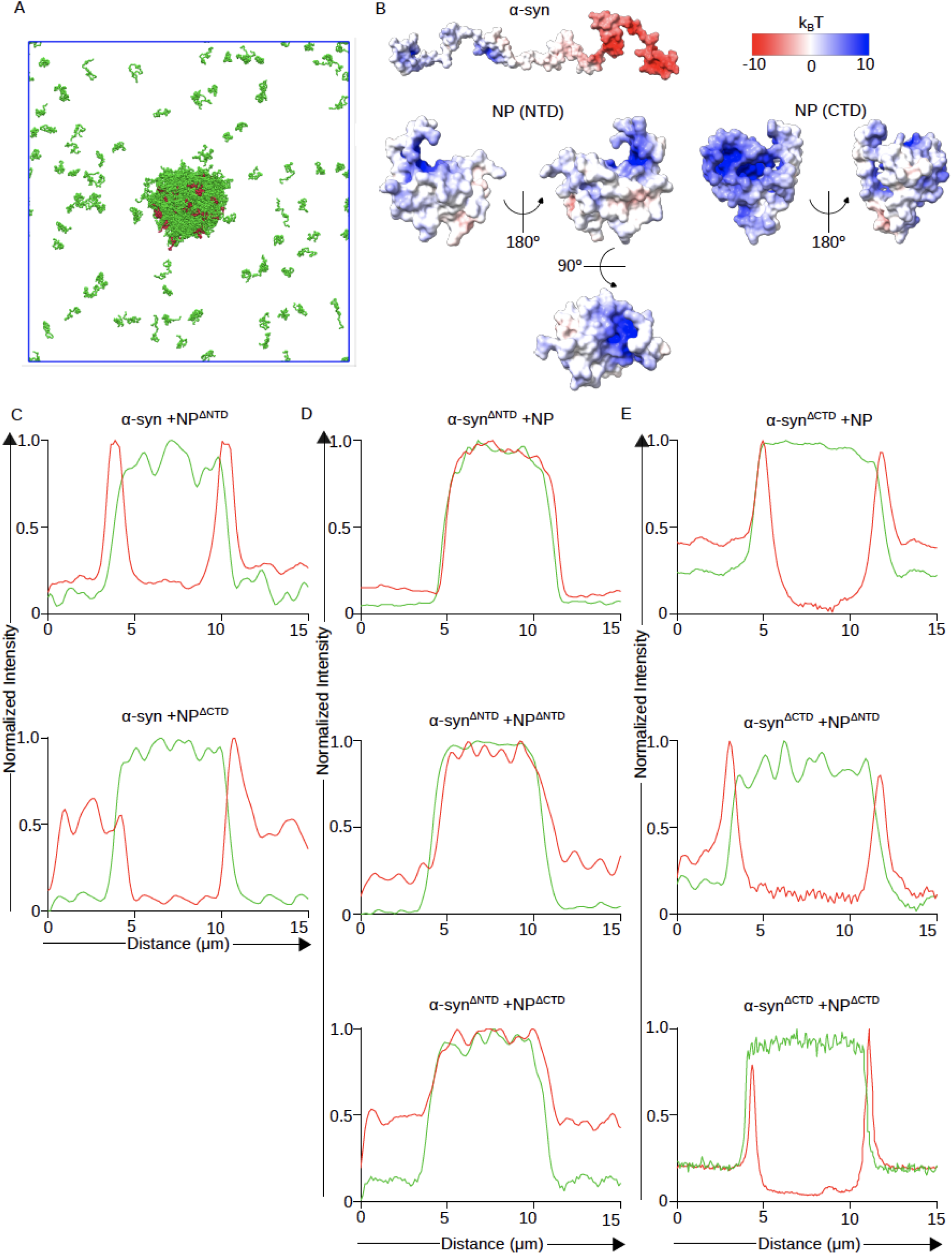
Coarse-grained (CG) simulation of α-syn and NP and line intensities of different co-condensates formed by different ɑ-syn and NP variants. (A) Simulation snapshot highlighting α-syn in green and NP in red in a cubic geometry. (B) Surface electrostatic potentials computed for the α-syn and of NP’s folded domain: NTD (PDB ID-7ACT) and CTD (PDB ID-6ZCO) showing the presence of distinct positive (blue) and negatively (red) charged surfaces. (C) Normalized intensity vs distance plot of ɑ-syn and NP truncated variants showing exclusion of NP truncated variants from the co-condensates. (D) Normalized intensity vs distance plot of ɑ-syn^ΔNTD^ and NP variants (WT and truncated) showing complete co-localization of ɑ-syn^ΔNTD^ and NP variants in the co-condensates. (E) Normalized intensity vs distance plot ɑ-syn^ΔCTD^ and NP variants (WT and truncated) showing exclusion of NP variants from the co-condensates.

**Figure S5:**
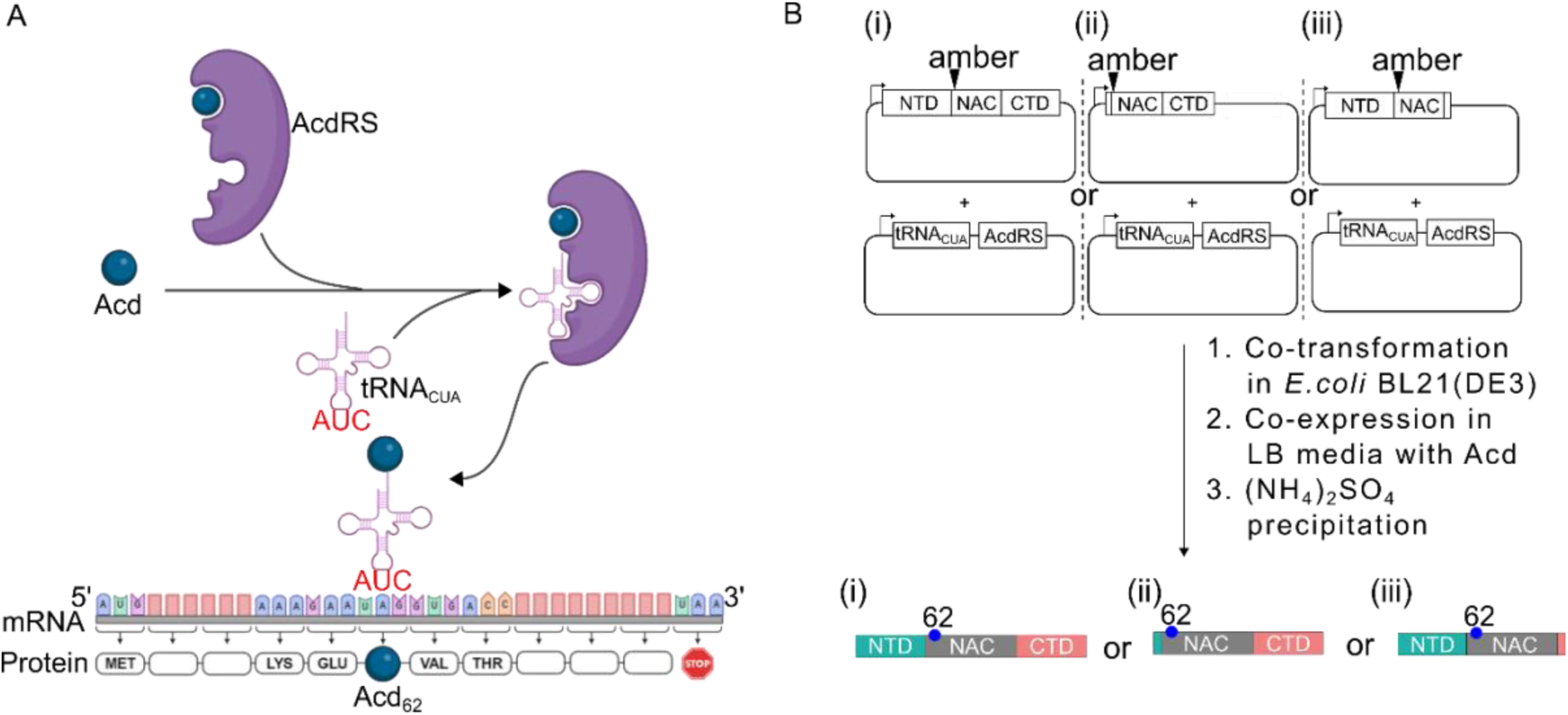
Schematic diagram showing the mechanism and workflow for site-specific incorporation of Acd in ɑ-syn. (A) Tyrosine aminoacyl tRNA synthetase (TyrRS) from *Methanococcus jannaschii* was strategically engineered through mutations (Y32G, L65E, D158G, and I158C) to create Acd aminoacyl tRNA synthetase (AcdRS), capable of incorporating Acd into its active site. The AcdRS enzyme charges tRNA_CUA_ with Acd, resulting in the formation of ^Acd^tRNA_CUA_. This modified tRNA recognizes the amber stop codon (TAG) and incorporates Acd at the corresponding position during translation. (B) To achieve site-specific incorporation of Acd, the genes encoding tRNA_CUA_ and AcdRS were co-transformed into *E. coli* using the pDule2 vector, along with the ɑ-syn-62TAG or its truncated constructs (ɑ-syn-62TAG^ΔNTD^ or ɑ-syn-62TAG^ΔCTD^) cloned in the pET-21A(+) vector. Chemically synthesized Acd was supplied in the growth medium while the *E. coli* cells were cultured for protein expression. Upon induction of the pDule2 vector, the AcdRS enzyme catalyzed the attachment of Acd to tRNA_CUA_, forming ^Acd^tRNA_CUA_. This charged tRNA recognized the TAG stop codon at the 62nd position of the ɑ-syn gene constructs. Consequently, Acd was site-specifically incorporated at the 62nd position of the WT and truncated variants (ɑ-syn^ΔNTD^ or ɑ-syn^ΔCTD^) of ɑ-syn protein during translation.

**Figure S6:**
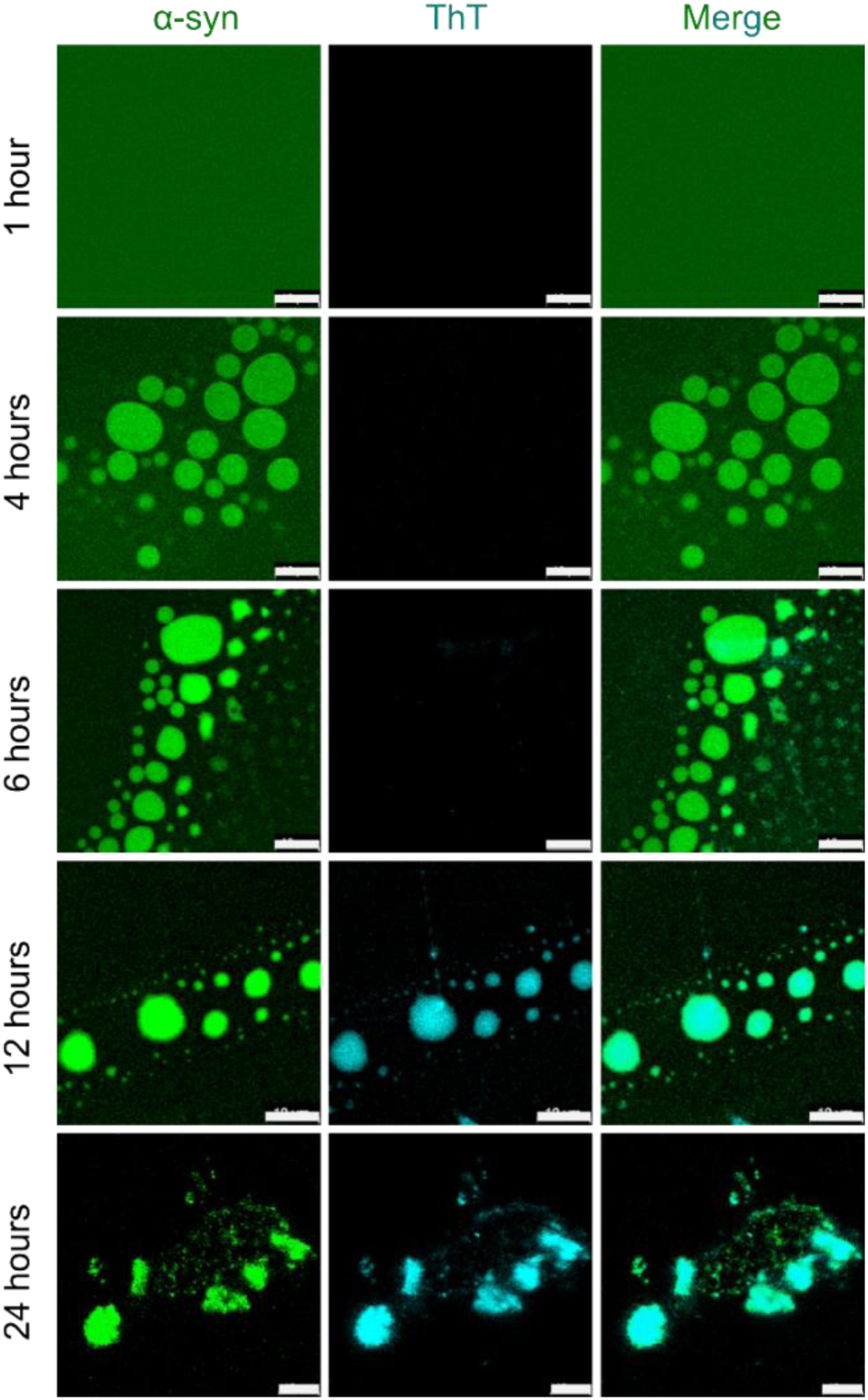
Temporal maturation of the ɑ-syn homotypic condensates. Time-dependent confocal microscopic images of 100µM ɑ-syn (doped with 1% Alexa-488 labeled ɑ-syn) co-incubated with 20µM ThT. The different channels are indicated at the top of the figure panel. The time points of incubation are indicated on the left of the figures. Scale bar: 10µm. Five independent experiments were done (n=5). The normalized ThT partition coefficients at each time points are plotted in Figure 5B.

**Figure S7:**
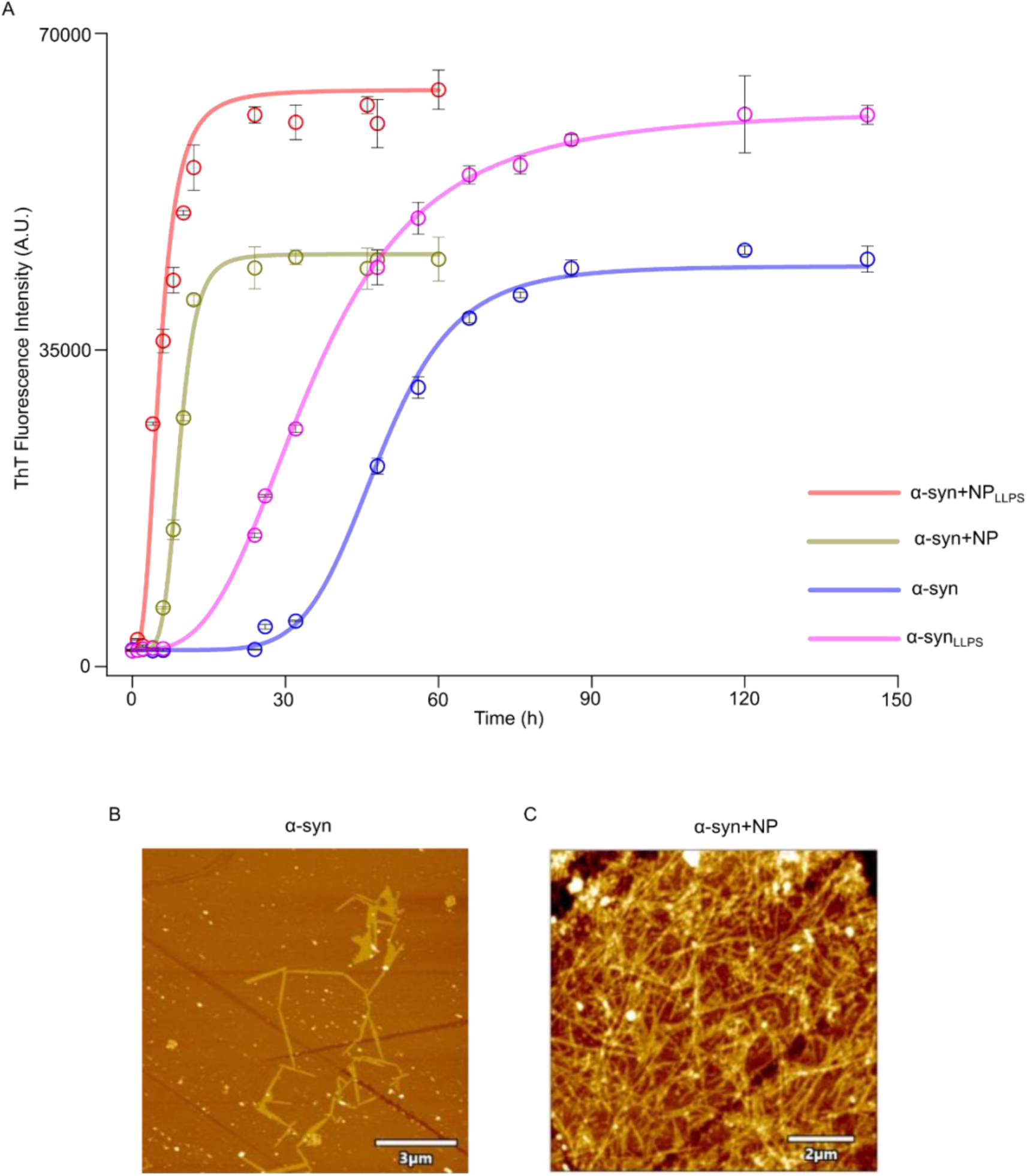
Modulation of ɑ-syn aggregation monitored by ThT fluorescence and AFM. (A) Plot of ThT fluorescence intensity vs time of incubation of different samples indicated in the legend. ɑ-syn and ɑ-syn_LLPS_ indicate 100µM molar protein in sodium phosphate buffer pH7.4 and 100µM protein in the presence of 10% PEG-8k (w/v), 400mM MgSO_4_ (LLPS condition) respectively. Whereas, ɑ-syn+NP and ɑ-syn+NP_LLPS_ indicate 100µM ɑ-syn+ 5µM NP co-incubated in sodium phosphate buffer pH7.4 and 100µM ɑ-syn + 5µM NP co-incubated in the presence of 10% PEG-8k (w/v), 400mM MgSO_4_ (LLPS condition) respectively. All the samples were kept at 37τC and 180 rpm shaking conditions. Three independent experiments were done with similar observations (n=3). (B) AFM images of ɑ-syn only fibril and fibril formed in the presence of NP. Three independent experiments were done with similar observations (n=3). The scale bar for (B) is 3µm and for (C) is 2µm. Three independent imaging were done with similar observations (n=3).

